# Modeling Human Spine-Spinal Cord Organogenesis by hPSC-Derived Neuromesodermal Progenitors

**DOI:** 10.1101/2023.07.20.549829

**Authors:** Dairui Li, Yuanchen Ma, Weijun Huang, Xiaoping Li, Huanyao Liu, Chuanfeng Xiong, Qi Zhao, Bin Wang, Xingqiang Lai, Shanshan Huang, Yili Wei, Junhua Chen, Xiyu Zhang, Lan Wei, Wenjin Ye, Qiumin Chen, Limin Rong, Andy Peng Xiang, Weiqiang Li

## Abstract

Human trunk development, including spine and spinal cord organogenesis, is a coordinated, orderly, and interdependent process with spatiotemporal tissue patterning. However, the underlying cellular and molecular mechanisms remain largely unclear due to the lack of an effective model that can simulate the early development of human body axis. Here, we reported the long-term patterning and dynamic morphogenesis of human trunk through the formation of spine-spinal cord organoids (SSCOs) self-organized from three-dimensional culture of human PSC-derived neuromesodermal progenitors (NMPs). The SSCOs resembled the morphogenetic features of spine and spinal cord along the anterior–posterior axis, and showed the chondro-osteogenic and neural trajectories consistent with developmental dynamics of spine and spinal cord in gestational embryo through single-cell RNA sequencing (scRNA-seq). In addition, we identified a new HMMR+ bipotent cell population with self-renewal ability and neural/mesodermal competence but distinct from NMPs, which may be involved in trunk development and represent an invaluable tool for disease modeling of spine- and spinal cord-related disorders.

**Graphic Abstract:** 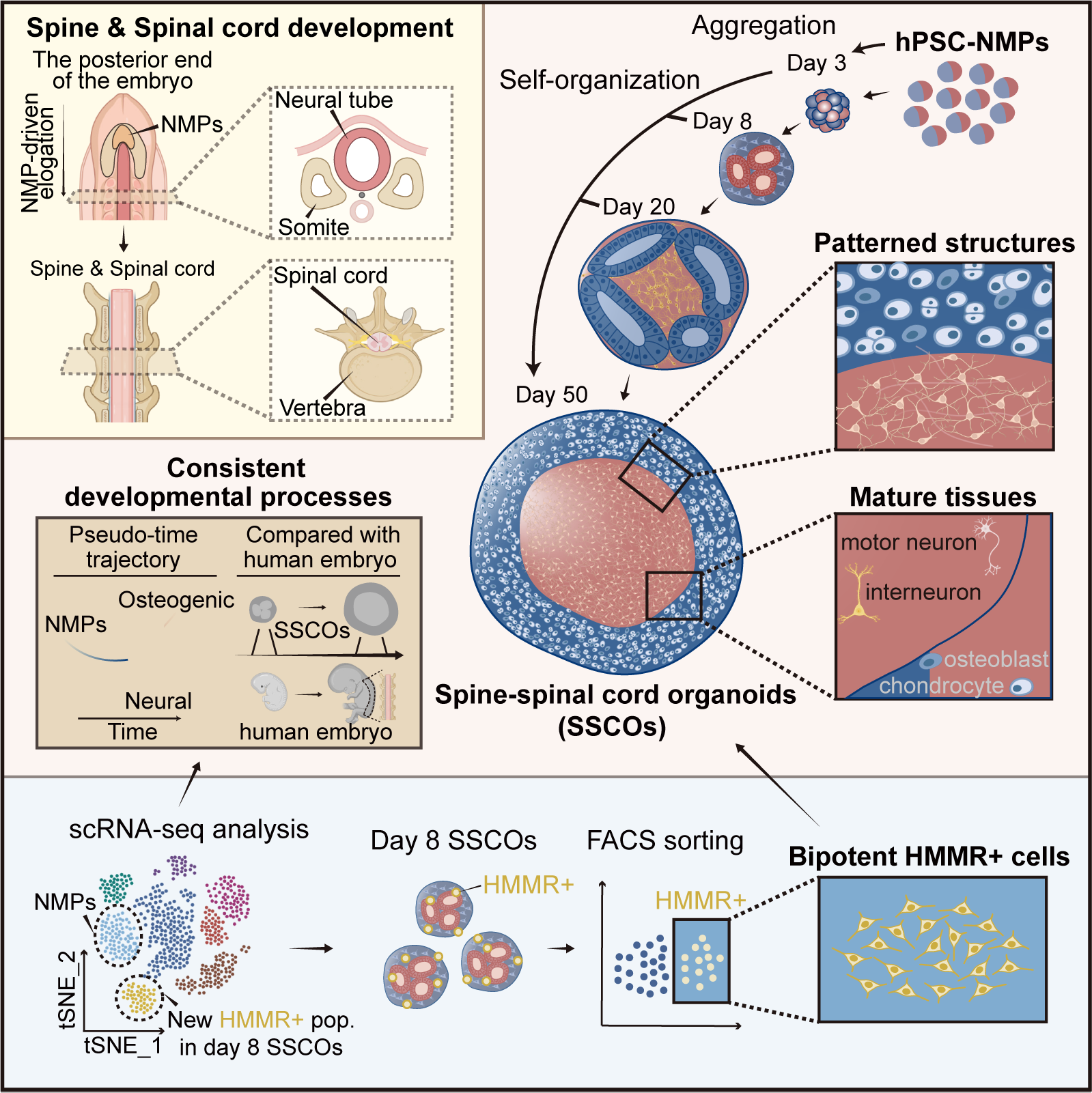

## Main

Early development of the spine and spinal cord starts with cell/tissue specialization. During the process of neurulation, epiblast cells in front of the anterior primitive streak are first specialized into the neural plate. The edges of the neural plate thicken and move upward to form the neural folds, and the neural folds are then closed to generate the neural tube, in which the anterior end develops into brain and the posterior portion becomes spinal cord. Posterior neural tubes are characterized by a “dorsal-ventral” axis, and motor neurons are mainly concentrated on the ventral side, while interneurons are in the rest of regions.^1^ The spine and its surrounding muscle tissues are developed from the paraxial mesoderm. During somitogenesis, the paraxial mesoderm differentiates into somites containing mesenchymal cells, which further gives rise to sclerotome containing chondroprogenitors. The sclerotome further encircles the sides of the neural tube and become precursors to the vertebra. Early vertebra precursors generate chondrification centers that gradually replace the sclerotome with cartilage. Primary and secondary ossification centers then arise in the cartilaginous vertebra and allow the vertebra to mature gradually.^2, 3^ Therefore, development of the spine and spinal cord is a complex multistep process and relies on the activity of different populations of axial progenitors.^4^ However, it is difficult to study their dynamics at the cellular and molecular level due to highly limited availability of human embryos under ethical and technical concerns. In recent years, three-dimensional (3D) organoid models were successfully developed for imitation of human development *in vitro*,^5^ which may provide an invaluable tool for studying human body axis elongation and spine/spinal cord organogenesis.

Despite belonging to the mesoderm and ectoderm respectively, retrospective clonal lineage analysis and lineage-tracing experiments revealed that the origin of the paraxial mesoderm and the posterior neural tube is not independent, but is derived from a special kind of axial progenitors: neuromesodermal progenitors (NMPs).^6, 7^ Therefore, NMPs play essential roles in development of the vertebrate trunk and may serve as an ideal cell model for exploring the developmental of human trunk and pathogenesis of trunk-related disorders. Jesse et al. achieved early mouse trunk-like structures (TLS) with neural tube and somites (corresponding to stage before E10.5) from mouse embryonic stem cells,^8^ while TLS with mature characteristics has not been addressed. Using pluripotent stem cell (PSC)-derived NMPs, Ju-Hyun et al. also generated functional human spinal cord organoids recapitulating neurulation-like tube-forming morphogenesis of the early spinal cord, rather than adjacent paraxial mesodermal tissues.^9^ Consequently, whether NMPs could form complex tissues containing spine and spinal cord simultaneously and how individual, neighboring components coordinate to establish a complex, functional trunk structures remains unknown.

Previous studies demonstrated that NMPs co-express definitive neural and mesodermal transcription factors, such as Sox2 and brachyury (Tbxt).^10, 11^ However, the NMPs may represent heterogeneous neuromesodermal competent cell (NMC) populations, containing some NMCs that actually give rise to both neural and mesodermal derivatives (referred to as NMPs), or other NMCs harboring both neural and mesodermal developmental potential but only generate either neural or mesodermal lineages.^4^ Moreover, NMPs are reported to emerge at the end of gastrulation at E7.5 but exhausted at the termination of axis elongation at E13.5.^12, 13^ Whether other types of neuromesodermal competent cells exist *in vivo* and contribute to long-term trunk tissue generation after NMP extinction remains to be clarified.

In this study, we reported the continuous patterning and dynamic morphogenesis of the spine-spinal cord organoids (SSCOs) as an easily accessible model from a three-dimensional culture of hPSC-NMPs. Single-cell RNA sequencing (scRNA-seq) was used to dissect the differentiation process and explore its relationship with in-vivo developmental trajectory in depth. The results indicated that SSCOs resembled the cooperative spine and spinal cord development and maturation. In addition, we also identified a new NMC-like population, bipotent HMMR+ cells, which originated from NMPs but showed different gene expression and localization patterns. Our results demonstrated that HMMR+ cells did not express SOX2 or TBXT, and could be propagated *in vitro* and differentiate to spinal cord motor neurons and mesenchymal cell lineages, which may be essentially involved in spine/spinal cord development and represent a potential cell model for elucidating the underlying molecular pathogenesis of spine- and spinal cord-related disorders.

## Results

### Efficient Generation and Characterization of hPSC-Derived NMPs

In this study, we first obtained NMPs from hPSCs (HEF-iPSC line)^14^ through activation of Wnt signaling (with CHIR99021), FGF signaling (with bFGF), and TGFβ signaling (with TGFβ1) for 2 days (Supplementary Fig. 1a,b) as previously reported.^14^ Immunostaining assay showed that most of the day 2 cells expressed NMP-specific genes including Brachyury (BRA) and SOX2 (Supplementary Fig. 1c). qPCR analysis revealed that day 2 cells expressed significantly lower levels of the pluripotency markers (*POU5F1, NANOG*) compared with day 0 cells, while the expression of NMP markers, such as *CDX1, WNT8A*, and *SP5* were highly upregulated after differentiation (Supplementary Fig. 1d). The single cell transcriptome data revealed that day 0 and day 2 cells were projected into 2 individual clusters after *t*-distributed stochastic neighbor embedding (*t*-SNE). Further analysis showed that the gene expression pattern of these 2 cell clusters was in accordance with qPCR results (Supplementary Fig. 1e-g). These results indicate that day 2 differentiated cells derived from hPSCs displayed similar gene expression patterns with NMPs, thus referred to as hPSC-NMPs.

To investigate the differentiation potential of hPSC-NMPs, day 2 cells were first induced into neural stem cells (NSCs) (from day 3 to day 6), and then differentiated into mature neurons for 4 weeks (from day 7 to day 34; Supplementary Fig. 2a). Neural rosettes resembling the developing neural tube were radially formed on day 6, containing cells expressing markers of NSCs such as SOX2, SOX1, PAX6, and NES (Supplementary Fig. 2b,c). In addition, expression of markers for neural progenitor cells (NPCs) such as *OLIG2* and *CDH6* were rapidly increased (Supplementary Fig. 2d). More importantly, motor neurons expressing TUJ1, ISL1, and ChAT could be detected on day 34, and neuronal markers such as *DCX, NEUROD1*, and *MAP2*, were highly upregulated compared with that of the cells on day 6 (Supplementary Fig. 2e,f). These results showed that hPSC-NMPs can differentiate into motor neurons through the intermediate stage of NSCs.

To further identify mesodermal differentiation potential of hPSC-NMPs, day 2 NMPs were cultured in NMP induction medium for additional 2 days (from day 3 to day 4), cells lost expression of SOX2 but were still strong positive for paraxial mesodermal progenitor (PMP) markers BRA and TBX6 (Supplementary Fig. 2g,h), indicating a transition from NMP to paraxial mesodermal state. Similar results were obtained by qPCR analysis since the expression of NMP markers (e.g., *WNT8A, FGF17, SP5, CDX1*, and *SOX2*) was downregulated obviously and the mRNA level of PMP markers (*BRA* and *TBX6*) was substantially increased (Supplementary Fig. 2i). When PMPs were cultured in MSC medium for 3 weeks (from day 5 to day 25), cells with mesenchymal cell morphology emerged (Supplementary Fig. 2j). These cells had similar expression pattern of cell surface markers and tri-lineage differentiation capacity with MSCs (Supplementary Fig. 2k-m). These results showed that hPSC-NMPs can generate mesodermal derivatives through PMPs.

The above evidence indicated the NMP cell population has the potential to differentiate to ectodermal and mesodermal lineages. However, whether a single human NMP has multipotency remains unclear. To verify this hypothesis, we performed clonal analysis and day 2 NMPs were dissociated and seeded as single cells at clonal density. We found that single NMPs could form colonies (Supplementary Fig. 2n,o), and about 30% of the colonies contained both TUJ1+ cells (neural lineage) and α-SMA+ smooth muscle cells (mesodermal lineage), indicating that single hPSC-NMPs are multipotent stem cells (Supplementary Fig. 2p).

### Self-organization of hPSC-derived NMPs in 3D culture

To investigate whether these cells could construct complex organoids containing both spine and spinal cord structures, hPSC-NMPs were first cultured in ultra-low attachment plates to form 3D aggregates on the shaker using N2B27 medium with growth factors (e.g., bFGF and EGF) for 6 days (from day 3 to day 8). Then 3D aggregates were allowed to proliferate and differentiate in medium containing neurotrophic factors (e.g., RA, BDNF, GDNF) and mesodermal lineage-inducing factors (e.g., BMP4, TGFB1) until day 20 (from day 9 to day 20) and mature in N2B27/chondrogenic mixed medium with RA and SAG until day 50 (Fig. 1a). During the differentiation process, the aggregates continued to expand (Fig. 1b,c), and the diameter of the aggregates increased obviously (∼200 μm on day 8; ∼700 μm on day 20; ∼1.5 mm on day 50) with a relatively low apoptosis rate (Fig. 1d and Extended Data Fig. 1a,b). To fully capture the molecular repertoire of subpopulation cells during ontogenesis of SSCOs at single-cell resolution, we conducted scRNA-seq of cells harvested from different stages using the BD Rhpasody platform (Fig. 1e) and obtained the characteristic gene expression profiles of each stage (Fig. 1f). HOX genes play essential roles in trunk morphogenesis along the anterior–posterior axis during vertebrate embryogenesis, which are clustered in four different clusters (HOXA, HOXB, HOXC, and HOXD) and consist up to 13 paralogous groups (PGs). Hox gene expression were first initialized in the posterior of the embryo, in a temporally progressive fashion that reflects their 3′-5′ genomic order (i.e. members of PG1 are activated first and PG13 last).^15-18^ We therefore analyzed the HOX gene expression pattern in SSCOs during differentiation process. We found that day 2 hPSC-NMPs highly expressed hindbrain HOX genes (1-3), and the cervical and thoracic HOX paralogs (4-9) were mildly expressed at this stage. Moreover, robust activation of HOX 4-9 was detected in differentiated cells on day 8-50, while 3’ HOX transcripts (rostral 1-3) remained expressed but with a relatively lower level. Interestingly, lumbosacral patterning (HOX10-13) was readily detected in differentiated cells on day 50. These results indicated the HOX gene expression pattern in SSCOs resembled the *in vivo* temporal features of HOX paralogs during the embryonic axis elongation, and caudal patterning may require prior activation of rostral HOX genes (Fig. 1g and Extended Data Fig. 1c).

**Fig. 1:**
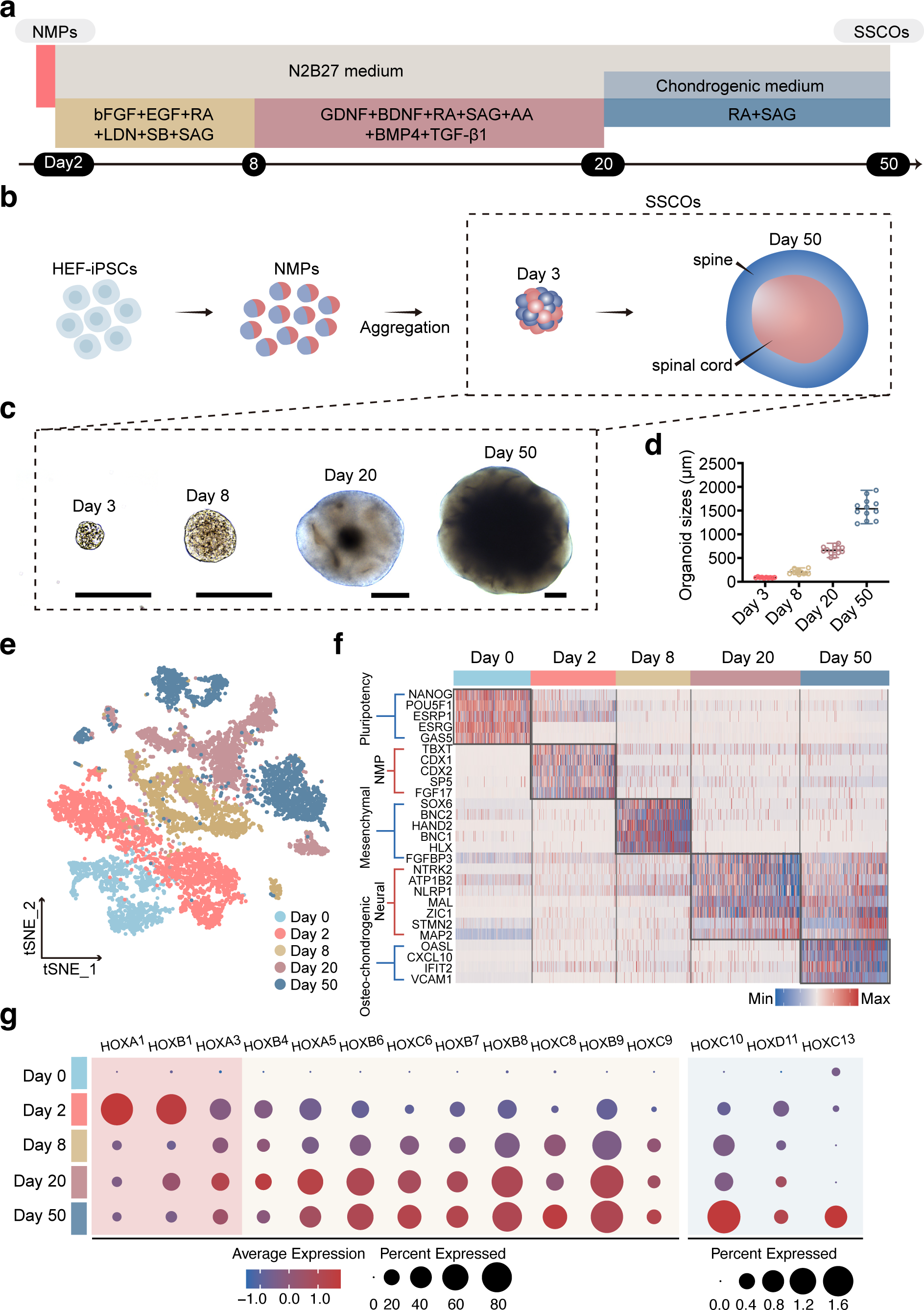
HEF-iPSC-derived NMPs Generate SSCOs in 3D Culture. **a,** Schematic of the strategy used to generate SSCOs from HEF-iPSC-derived NMPs. **b,** Schematic overview for generation of SSCOs from HEF-iPSC-derived NMPs. **c,** Bright-field images of SSCOs at different stages (days 3, 8, 20 and 50). Scale bar, 250 μm. **d,** Graph showing sizes of SSCOs (average diameter) during different development stages. n = 10 (day 3), 12 (day 8), 12 (day 20), 12 (day 50). **e,** *t*-SNE plot of SSCOs under different developmental period, colored with different developmental stages. **f,** Heatmap of stage-enriched genes. Each column represents a single cell and each row represents one characteristic gene. The colors ranging from blue to red indicate low to high relative gene expression levels. **g,** Dot plot of stage-enriched HOX genes. The dot color represents the average expression level of selected genes and dot size represents the percent expressed cells in each cluster.

On Day 8, the aggregates of hPSC-NMPs formed round-shaped structures with distinct edges (Day 8 SSCOs; Fig. 2a). Immunofluorescence staining showed the evidence of lineage segregation in Day 8 SSCOs, since we could detect a region corresponding to neuroectoderm expressing SOX2, PAX2 and NES, and a mesodermal region, as evidenced by SOX9, PGDFRA and PDGFRB expression. Notably, expression of SOX10 was not detected in day 8 SSCOs, suggesting the absence of neural crest cells in these aggregates (Fig. 2b and Extended Data Fig. 1d). qRT-PCR revealed that the expression of NMP markers (e.g., *BRA, TBX6*, and *MIXL1*) were greatly reduced and markers indicating neuroepithelial cells (e.g., *SOX2, PAX2, CDH6*, and *PAX6*) and mesenchyme progenitors with somitic markers (e.g., *FOXC2, NKX3-2, SOX9*, and *MEOX2*)^3^ were substantially increased compared with day 2 NMPs (Fig. 2c). In addition to the three major cell types including neural stem cells (NSCs; expressing *SOX2, TOP2A* and *ZIC1*), interneuron progenitors (INPs; expressing *PAX2, LHX1*), and mesenchymal progenitors (MPs; expressing *SOX6, NKX3-2, PDGFRA*, and *TBX18)* presented in day 8 SSCOs, we also observed endothelial progenitor cells (EPCs; expressing *CD34, ESAM*) and a new population (New pop.) within day 8 SSCOs scRNA-seq data. This new population exhibited co-expression of markers associated with NSCs (*TOP2A, CENPF*) as well as MPs (*PDGFRA, NKX3-2* and *SOX6*) (Fig. 2d-f). Based on these data, day 8 SSCOs were identified as the “somite-neuroepithelium genesis” stage.

**Fig. 2:**
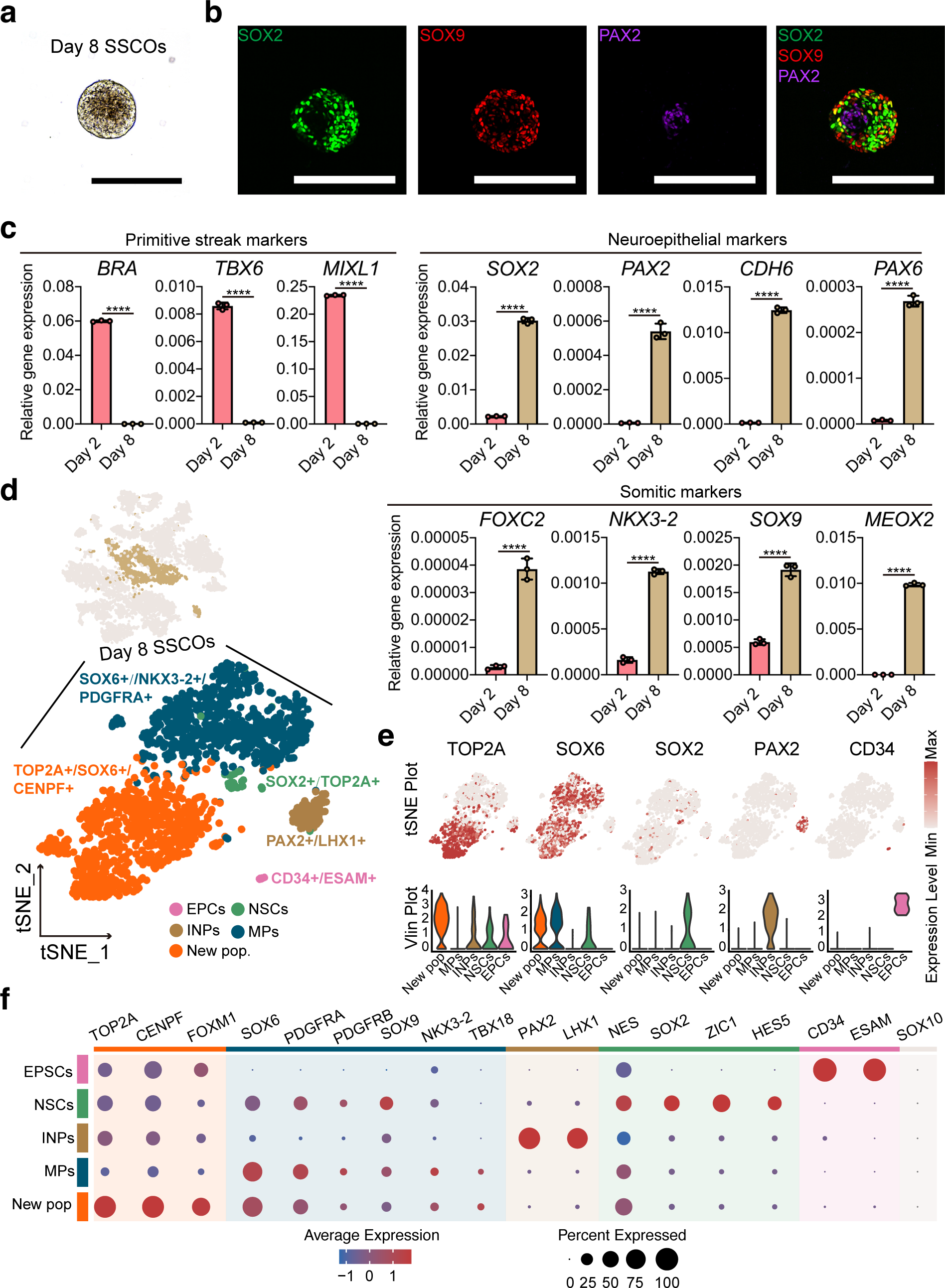
NMPs Self-organize to Day 8 SSCOs Containing Mesenchymal and Neural Stem Cells. **a,** Bright-field images of SSCOs at day 8. Scale bar, 250 μm. **b,** Immunofluorescence analysis of sectioned day 8 SSCOs. Scale bar, 250 μm. **c,** qPCR analysis for the gene expression of primitive streak markers, neuroepithelial markers, and mesenchymal markers in day 8 SSCOs compared with day 2 NMPs. Data are represented as mean ± SD. Unpaired two-tailed t test is used for comparison of two groups. *p < 0.05, **p < 0.01, ***p < 0.001, ****p < 0.0001. **d,** *t*-SNE plot of scRNA-seq profiles showing clusters of neural stem cells (NSCs), mesenchymal progenitors (MPs), interneuron progenitors (INPs), endothelial progenitor cells (EPCs) and a new population (New pop.) in day 8 SSCOs. **e,** Selected gene expression of mixed cluster in *t*-SNE plot (upper panel) and violin plot (lower panel), in *t*-SNE plot each genes distribution and relative expression levels were scaled from grey (low expression) to red (high expression), and in violin plot the y-axis represents the relative expression level of selected genes. **f,** Dot plot of cluster-enriched genes clusters in day 8 SSCOs. The dot color represents the average expression level of selected genes and dot size represents the percent expressed cells in each cluster.

To further explore interactions among the new population, NSCs and MPs, we performed cell cross talking analysis in day 8 SSCOs with CellChat software.^19^ The results indicated that both MPs and new population revealed a higher level of cross talking ability compared with NSCs (Extended Data Fig. 2a-c). Further signaling transduction exploration pointed that these cell clusters could be separated into three communication patterns (Extended Data Fig. 2d,e). Notably, the new population received LAMNIN (LAMA5-DAG1) and AGRN (AGRN-DAG1) signalings from MPs (Extended Data Fig. 2f). The new population could also interact with NSCs through JAM (JAM3-JAM3) signaling pathway, and regulate both MPs and NSCs through PTN signaling pathway with different ligand-receptor pairs (PTN-NCL and PTN-SDC3) (Extended Data Fig. 2f). The function of these signaling pathways during SSCO development needs to be further clarified.

### Neural and mesenchymal lineages development in SSCOs

Day 8 SSCOs were further cultured in suspension in medium supplemented with both neurotrophic and mesodermal lineage-inducing factors^20-26^ including GDNF, BDNF, AA, RA, SAG, BMP4 and TGF-β1 for 12 days. Through continuous induction, tubular structures appeared at the edge of the day 20 SSCOs (Fig. 3a), which was the region of chondroprogenitors (CPs) expressing SOX9, while cells in the central region of SSCOs expressed the neuronal markers TUJ1 and DCX, indicating further differentiation and morphogenesis of SSCOs. (Fig. 3b and Extended Data Fig. 1e). In addition, immunofluorescence staining showed a slightly larger cartilage area (SOX9+; 50-60%) than neural area (TUJ1+; 40-50%; Extended Data Fig. 1f). Compared with day 8 SSCOs, expression levels of some mesenchymal and neuroepithelial transcription factors were reduced (e.g., *MEOX2, TWIST1*) or remained unchanged (e.g., *FOXC2, CDH6*) (Fig. 3c), while expression of neural lineage including neural progenitor cells (*PAX6*) and immature neurons (INs) (e.g., *DCX, ISL1*, and *NEFL*) markers, and chondrogenic markers (e.g., *MSX1, SOX9, IBSP*, and *ACAN*) were markedly increased (Fig. 3c). Three cell clusters were profiled in day 20 SSCOs by scRNA-seq, defined as neural progenitor cells (NPCs, expressing *FABP7, MSI1* and *NES*), INs (expressing *DCX, NHLH1* and *ELAVL3*), and CPs (expressing *MGP, POSTN* and *FOXC1*) (Fig. 3d-f). In addition, dorsal-ventral patterning of both neural tube and sclerotome were detected at this day 20 SSCOs stage according to the pattern diagram (Fig. 3g)^3, 27-29^. It seemed that dorsal genes (e.g., *LMX1A, MSX1, MSX2, OLIG3, PAX3*, and *PAX7*) were more abundant compared with the ventral markers (e.g., *IRX3, FOXN4, NKX6-1, OLIG2*, and *LMX1B*) in NSCs and INs of day 20 SSCOs (Fig. 3h)^30-32^. Dorsal markers (e.g., *MSX1, MSX2*), central/ventral markers (e.g., *PAX9, NKX3-2*), lateral markers (e.g., *KDR, SIM1*) of sclerotome and the marker of syndetome (SCX) were also detected in CPs of day 20 SSCOs (Fig. 3h). Moreover, with immunofluorescence staining, we found that day 20 SSCOs contained OLIG2+ or ISL1+ motor neuron progenitors, ChAT+/HOXC6+ motor neurons, and PAX2+ interneuron progenitors (Fig. 3i and Extended Data Fig. 1g). Therefore, day 20 SSCOs were indicated as “sclerotome-neurogenesis” stage. These results showed that SSCOs were able to develop into more mature structures.

**Fig. 3:**
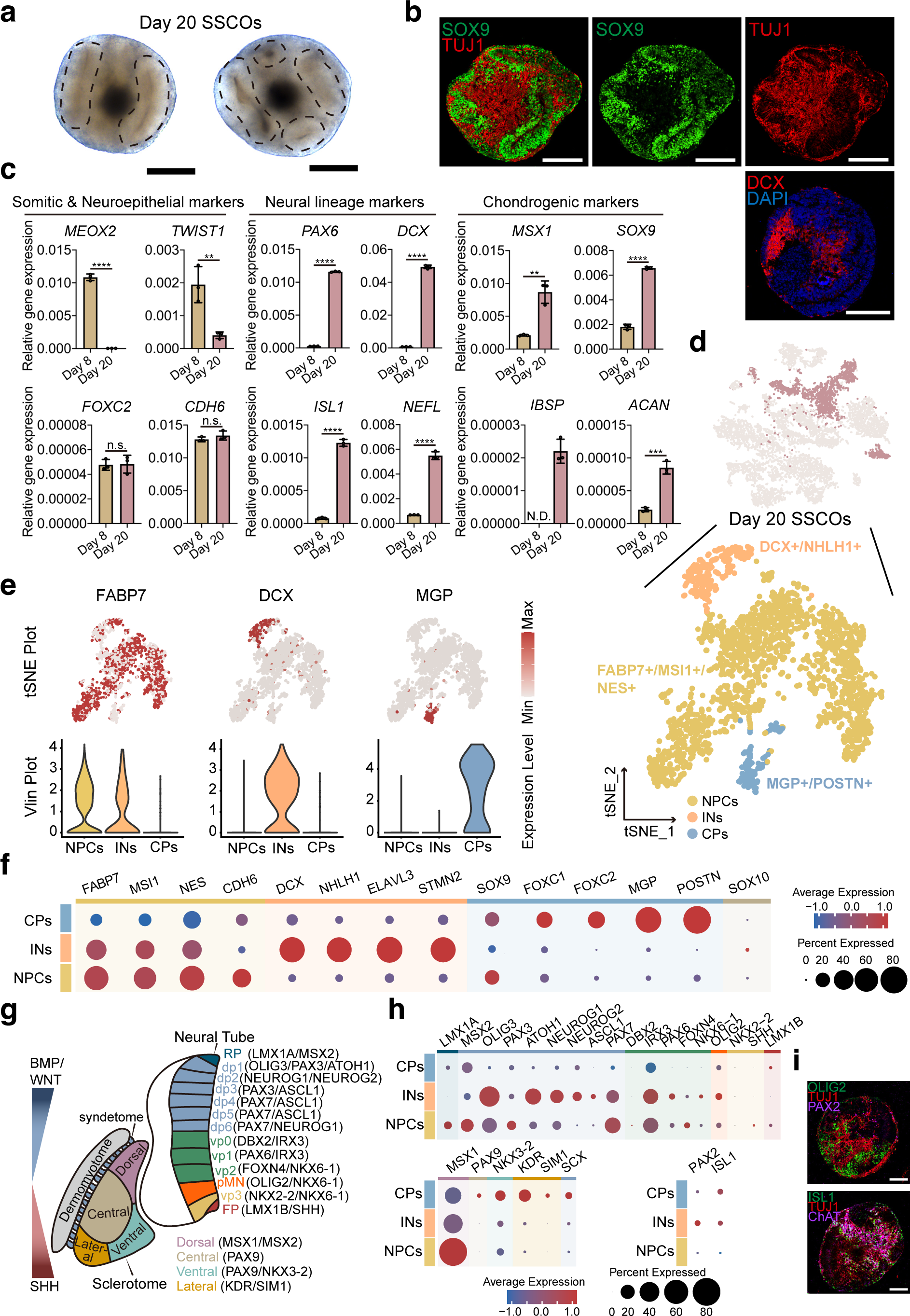
Chondroprogenitors and Immature Neurons Appear in Day 20 SSCOs. **a,** Bright-field images of SSCOs at day 20. Scale bar, 250 μm. **b,** Immunofluorescence analysis of sectioned day 20 SSCOs showing the expression of the chondroprogenitor marker SOX9 and immature neuron marker TUJ1, DCX. Scale bar, 250 μm. **c,** qPCR analysis of neuroepithelial & somitic markers, neural markers and chondrogenic markers in day 20 SSCOs compared with day 8 SSCOs. Data are represented as mean ± SD. Unpaired two-tailed t test is used for comparison of two groups. n.s., not significant. N.D., not detected. *p < 0.05, **p < 0.01, ***p < 0.001, ****p < 0.0001. **d,** *t*-SNE plot of scRNA-seq profiles showing clusters of neural progenitor cells (NPCs), immature neurons (INs) and chondroprogenitors (CPs) in day 20 SSCOs. **e,** Gene expression of representative markers among NPCs, INs and CPs in *t*-SNE plot (upper panel) and violin plot (lower panel), in *t*-SNE plot each genes distribution and relative expression level was scaled from grey (low expression) to red (high expression), and in violin plot the y-axis represents the relative expression level of selected genes. **f,** Dot plot of cluster-enriched genes of clusters in day 20 SSCOs. The dot color represents the average expression level of selected genes and dot size represents the percent expressed cells in each cluster. **g,** Schematic of dorsal-ventral patterning of neural tube and sclerotome. **h,** Dot plot of genes showing dorsal-ventral patterning of day 20 SSCOs. **i,** Immunofluorescence analysis of sectioned day 20 SSCOs showing the expression of the motor neuron progenitor marker OLIG2, ISL1 and the interneuron progenitor marker PAX2. Scale bar, 150 μm.

The orderly development and precise arrangement of the spine and spinal cord requires the synergy of multiple signaling pathways. To further explore whether there are interactions between mesodermal derivatives and neural derivatives during organoid development, we also performed cell cross talking analysis in in day 20 SSCOs. The result indicated that CPs and NSCs both revealed a higher level of cross talking ability compared with INs (Extended Data Fig. 2g-i). Further signaling transduction exploration indicated that each cell cluster could be completely separated into different communication patterns (Extended Data Fig. 2j,k). Among these patterns, we found that CDH (CDH2-CDH2), CADM (CADM3-CADM3) and FN1 (FN1-ITGA5) signaling pathways were mainly involved in self-regulation of NPCs, INs and CPs respectively (Extended Data Fig. 2i,l), which was similar with previous reports.^33-35^ Remarkably, ncWNT signaling, which was essential for early spinal cord development,^36, 37^ mainly targeted to NSCs and INs in our results. Interestingly, ncWNT ligands were predominantly secreted from CPs in SSCOs, which has not been reported (Extended Data Fig. 2l). NCAM signaling mainly contributed to neurogenesis and was also implicated in skeletogenesis and chondrogenesis.^38-40^ In our results, molecules of NCAM signaling targeted to CPs were primarily derived from INs (Extended Data Fig. 2l). IGF signaling was involved in chondrogenesis in according to the previous studies.^41, 42^ Interestingly, NPCs and INs respectively secreted IGF2 to co-regulate the development of CPs (Extended Data Fig. 2l). The above evidence indicated that the interaction between neural derivatives and the adjacent mesoderm lineages may rely on diverse signaling pathways during early trunk development.

### Maturation of spinal cord neurons and vertebral chondrocytes and osteoblasts in SSCOs

To promote the rostrocaudal elongation and maturation of neurons and chondrocytes in organoids, day 20 SSCOs were cultured in chondrogenic medium supplemented with neural induction factors RA (for posteriorisation) and SAG (for ventralisation) for additional 30 days (Fig. 4a). Distinct structures in the lateral and the medial parts of day 50 SSCOs were formed as shown by HE staining and toluidine blue staining (Fig. 4b). Complex but finely organized tissues containing the cartilage region expressing COL-2 and CTSB surrounding the neural region expressing TUJ1 and MAP2 (Fig. 4c) was detected. Calcium imaging analysis using Fluo-4 acetoxymethyl ester (Fluo-4 AM) staining indicated that most of the neurons displayed spontaneous intracellular calcium oscillations (Extended Data Fig. 3a and Supplementary Video 1), and whole-cell patch-clamp analysis showed that some of neurons could fire sustained action potential (Extended Data Fig. 3b). In addition, compared with day 20 SSCOs, day 50 SSCOs highly expressed the markers of mature neurons (MNs) (e.g., *LBX1, HB9*), chondrocytes (CCs) (e.g., *COL-2, IBSP*, and *ACAN*), and osteoblasts (OBs) (e.g., *RUNX2, COL-1, ALP*, and *OPN*) (Fig. 4d). Four cell clusters were profiled in day 50 SSCOs by scRNA-seq, defined as INs (expressing *DCX, NCAM1*), MNs (expressing *MAP2, GAP43*), CCs (expressing *SOX9, VEGFA*), and OBs (expressing *COL1A1, COL1A2*) (Fig. 4e-g). Further, we showed that INs and MNs in day 50 SSCOs displayed the dorsal/ventral identity through analyzing the expression of dorsal or ventral specific markers (Fig. 4h,i)^27, 28^, which showed stronger expression of ventral genes than dorsal markers. We also identified ventral HB9+ motor neurons and dorsal LHX1+ interneurons with immunofluorescence staining in day 50 SSCOs (Fig. 4j). Therefore, day 50 SSCOs were recognized as “cartilage-neuron maturation” stage.

**Fig. 4:**
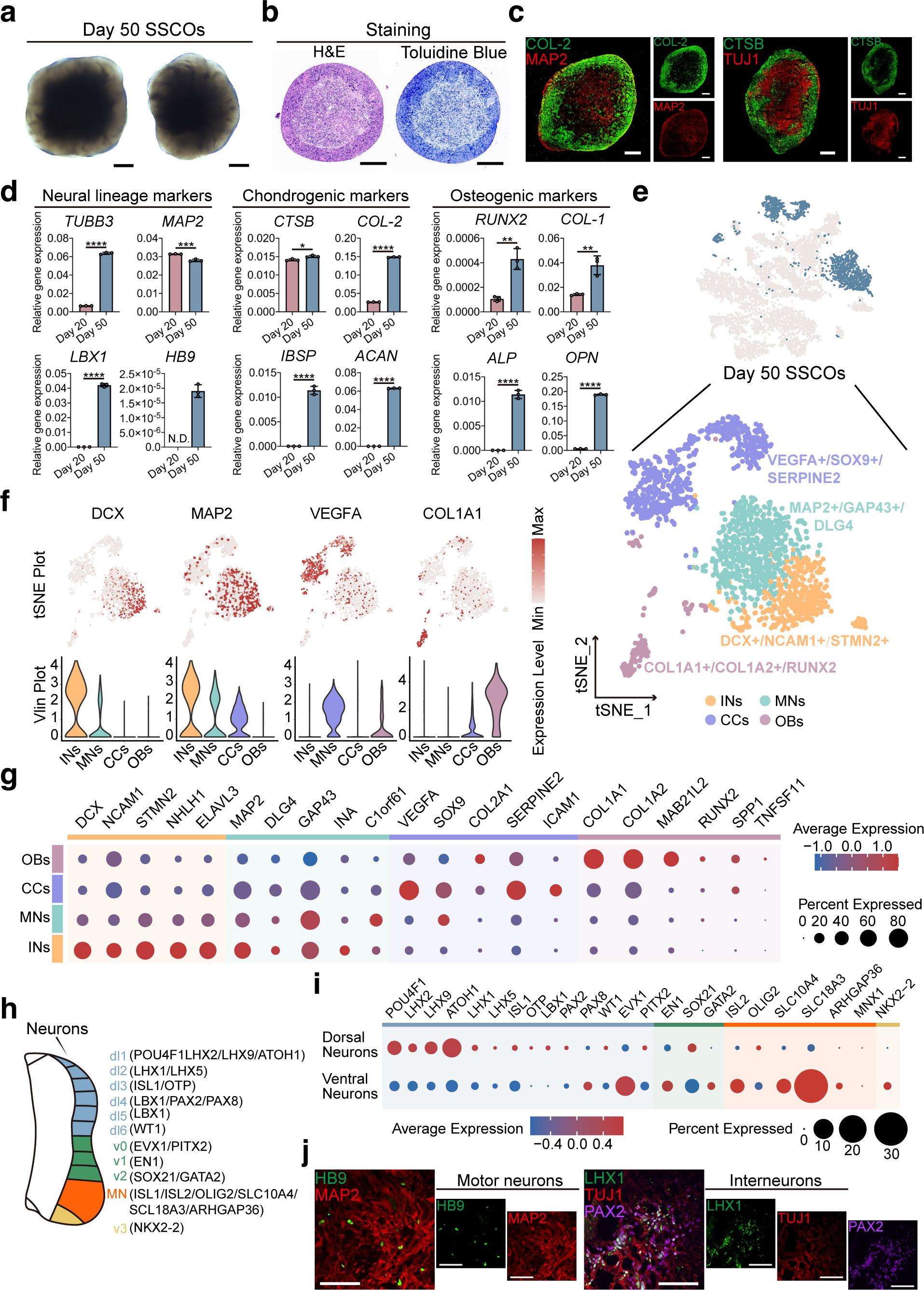
Self-Organization of Patterned Structures with Mature Spinal Cord Neurons and Chondrocytes in Day 50 SSCOs. **a,** Bright-field images of SSCOs at day 50. Scale bar, 250 μm. **b,** H&E staining and Toluidine Blue analysis of sectioned day 50 SSCOs. Scale bar, 250 μm. **c,** Immunofluorescence analysis of sectioned day 50 SSCOs showing the expression of the chondrocyte markers (COL-2, CTSB) and mature neuron marker MAP2. Scale bar, 250 μm. **d,** qPCR showing the expression of neural, chondrogenic and osteogenic markers in day 50 SSCOs compared with day 20 SSCOs. Data are represented as mean ± SD. Unpaired two-tailed t test is used for comparison of two groups. N.D., not detected. *p < 0.05, **p < 0.01, ***p < 0.001, ****p < 0.0001. **e,** *t*-SNE plot of scRNA-seq profiles showing clusters of immature neurons (INs), mature neurons (MNs), chondrocytes (CCs) and osteoblasts (OBs) in day 20 SSCOs. **f,** Gene expression of representative markers among INs, MNs, CCs and OBs in *t*-SNE plot (upper panel) and violin plot (lower panel), in *t*-SNE plot each genes distribution and relative expression was scaled from grey (low expression) to red (high expression), and in violin plot the y-axis represents the relative expression of selected genes. **g,** Dot of cluster-enriched genes of clusters in day 50 SSCOs. The dot color represents the average expression level of selected genes and dot size represents the percent expressed cells in each cluster. **h,** Schematic of dorsal-ventral patterning of spinal cord neurons. **i,** Dot plot of genes showing dorsal-ventral patterning of neurons in day 50 SSCOs. The dot color represents the average expression level of selected genes and dot size represents the percent expressed cells in each cluster. **j,** Immunofluorescence analysis of sectioned day 50 SSCOs showing the expression of the motor neuron marker HB9 and interneuron markers PAX2 and LHX1. Scale bar, 100 μm.

We performed cell cross talking analysis in day 50 SSCOs. The result indicated that both OBs and CCs revealed a higher level of cross talking ability compared with INs and MNs (Extended Data Fig. 2m-o). Further signaling transduction exploration pointed that these cell clusters could be separated into two outgoing and three incoming communication patterns (Extended Data Fig. 2p,q). Among these patterns, CCs and OBs not only showed strong self-regulation but also had active connections with other clusters (Extended Data Fig. 2n). Selected significant ligand-receptor pairs analysis revealed a panel of active signalings like COLLAGEN, BMP and FGF, which was vital for chondrogenesis or osteogenesis (Extended Data Fig. 2o).^43-45^ Apart from the above mentioned BMP, FGF and COLLAGEN pathways, we also found EPHA (ENNA3-EPHA7) signaling, which was known for axon guidance and neuron development^46^, was specifically secreted from CCs (Extended Data Fig. 2r).

Moreover, to promote further development and maturation of SSCOs, day 30 samples were transplanted into the kidney capsule of NCG mice (Extended Data Fig. 3c). We found that the transplanted organoids could survive and continue to grow in almost all mice 8 weeks after transplantation (Extended Data Fig. 3d). Samples were further dissected and analyzed by histochemistry assays. Cartilaginous tissues with characteristic chondrocyte morphology encircling neural-like tissues were identified by H&E staining and then validated by Safranin O staining and toluidine blue staining (Extended Data Fig. 3e). Immunofluorescence staining revealed that these tissues contained both intensely stained cartilaginous region expressing ACAN and neuronal region expressing MAP2 (Extended Data Fig. 3f), while the existence of human nuclei (HuNu)-positive cells in these regions indicated that transplanted tissues were indeed derivatives from human cells (Extended Data Fig. 3f). These results demonstrated that SSCOs also can differentiate and mature *in vivo*.

To verify the reproducibility of our organoid differentiation protocol, we also used two other PSC lines (PBMC-derived hiPSC line, H9 human ESC line) to generate SSCOs, and immunofluorescence staining identified that two key populations, including mesenchymal and neural lineages with characteristic distributions existed in the organoids during differentiation process (Extended Data Fig. 4a,b).

### SSCOs model the *in vivo* development of human spine and spinal cord

So far, we identified 13 cell clusters including hPSCs, NMPs, New pop., EPCs, NSCs, INPs, NPCs, INs, MNs, MPs, CPs, CCs, and OBs in our scRNA-seq data (Fig. 5a). Pseudo temporal analysis revealed two trajectories rooted in the NMPs and branched at day 8 by the “DDRTree” method (Fig. 5b), one of which was osteogenic trajectory in sequence as “NMPs-New pop-CPs-CCs-OBs”, while the other was neural trajectory in sequence as “NMPs-New pop-NPCs-INs-MNs”, further suggesting that the SSCOs recapitulates the developmental dynamics of spine and spinal cord in gestational embryo (Fig. 5b).

**Fig. 5:**
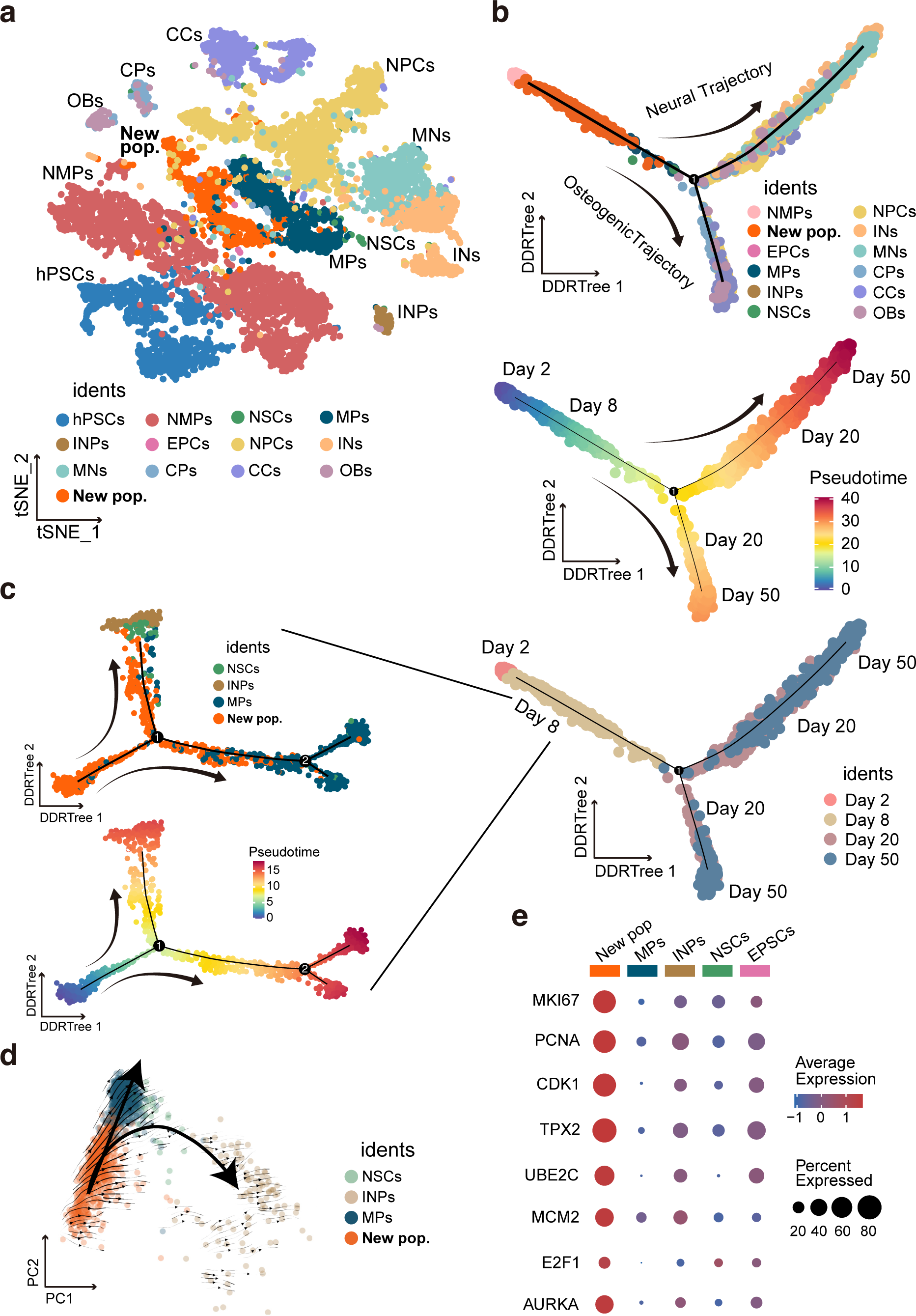
scRNA-Seq Analysis Reveals a New Population in Day 8 SSCOs. **a,** *t*-SNE plot of scRNA-seq profiles, colored with annotated cell clusters. **b,** Pseudotemporal analysis shows neural and osteogenic developmental trajectories in SSCO model initiated from NMPs and branched at day 8 stage. The colors denote cell types in the upper panel. Pseudotime is scaled from blue to red in the middle panel. Stages are denoted in the lower panel. **c,** Pseudotemporal analysis of day 8 SSCOs shows the new population could give rise to both mesenchymal progenitors and neural stem cells. The colors denote cell types in the upper panel. Pseudotime is scaled from blue to red in the lowe panel. **d,** RNA velocity analysis visualized in PCA plot shows neural and mesenchymal developmental trajectories in day 8 SSCOs initiated from the new population. **e,** The dot plot shows the new population expresses proliferative markers. The dot color represents the average expression level of selected genes and dot size represents the percent expressed cells in each cluster.

To further explore whether the *in vitro* SSCO differentiation model is consistent with spine-spinal cord organogenesis *in vivo*, we compared our scRNA-seq data of SSCOs to published human embryo scRNA-seq data from GW2-2.5 whole embryo (E-MTAB-9388)^47^, GW4-7 spinal cord (GEO: GSE171892)^48^, and GW8-25 spinal cord (GEO: GSE136719)^49^ (Extended Data Fig. 5a). The results revealed that most cells in day 8 SSCOs could overlap with the *in vivo* data (Extended Data Fig. 5b). We found that NSCs in day 8 SSCOs were mainly overlapped with GW6-7 spinal cord samples according to the expression of neuroectodermal markers SOX2 and HES5 (Extended Data Fig. 5c,d). Furthermore, 12.2% of MPs and 82.7% of new population cells in day 8 SSCOs were overlapped with GW4-5 spinal cord samples, while 67.3% of MPs and 6.0% of new population cells were overlapped with GW6-7 spinal cord samples according to the expression of mesodermal markers FOXC1 and TWIST1 (Extended Data Fig. 5c,d). NPCs and INs in day 20 SSCOs were also overlapped with *in vivo* data (Extended Data Fig. 5e). 97.4% of INs was mainly overlapped with GW7-17 spinal cord samples with the expression of neuronal markers DCX and STMN2 (Extended Data Fig. 5f,g). 34.2% of FABP7+ NPCs was overlapped with all GW7-25 spinal cord samples (Extended Data Fig. 5f,g). In day 50 SSCOs, 88.1% of INs (defined with markers DCX/STMN2) and 90.8% of MNs (defined with markers MAP2/GAP43) were mainly overlapped with GW7-17 spinal cord samples, with a small amount of INs and 8.2% of MNs were correlated with all GW7-25 spinal cord samples (Extended Data Fig. 5h-j). In summary, these results suggested that the *in vitro* differentiation of SSCOs could simulate the developmental process of GW4-25 spinal cord *in vivo*.

### Identification of a new bipotent HMMR+ cell population with neural and mesodermal competence in SSCOs

Although our results revealed that osteogenic and neural trajectory during SSCO formation. However, it has branched at day 8 clusters instead of day 2 NMPs by pseudo temporal analysis (Fig. 5b). To detailedly describe this new population, we analyzed day 8 SSCOs separately by pseudo temporal analysis, and found that the new population could give rise to both mesenchymal progenitors and neural stem cells (Fig. 5c). Moreover, RNA velocity^50^ analysis showed that neural and mesenchymal developmental trajectories in day 8 SSCOs initiated from the new population (Fig. 5d). Both of these results indicated that MPs, NSCs and INPs were derived from the new population, indicating that this population may represent a new bipotent cell progenitors with neural and mesodermal differentiation potential distinct from NMPs. Proliferative markers (e.g., *MKI67, PCNA, CDK1* and *TPX2*; fig. 5e) were significantly expressed in the new population, suggesting that it may play an active role during trunk development.

We further revealed that this new population (SOX6+/TOP2A+ cells) specifically expressed the cell surface marker *HMMR* (CD168) by interpreting the scRNA-seq data (Fig. 6a). Then we sorted HMMR+ cells in day 8 SSCOs through FACS, with the ratio of about 1-3% (Fig. 6b), similar to the ratio of that in GW2-2.5 and GW7-25 spinal cord samples (Extended Data Fig. 6c). HMMR+ cells can proliferate *in vitro* and be passaged at least 10 times while retaining HMMR and TOP2A expression (Fig. 6c,d). To verify the differentiation potentials of HMMR+ cells, we first seeded HMMR+ cells and HMMR-cells after FACS as single cells at a very low density. We found that single HMMR+ cells could grow into colonies, which contained cells expressing neural marker TUJ1 or mesodermal marker α-SMA, while only α-SMA+ cells could be detected in HMMR-cell-derived colonies (Fig. 6e,f). About 24.17%±5.34 of individual CD168+ cells derived from day 8 SSCOs (HEF-iPSC line) could form colonies. Moreover, we found that 66.65% ± 3.17% of the colonies formed by individual CD168+ cells contained both have TUJ1+ neural cells and α-SMA+ mesodermal cells in three batches using HEF-iPSCs (Fig. 6g). To further verify whether HMMR+ cells can generate SSCOs, HMMR+ cells were cultured in suspension to form 3D aggregates in ultra-low attachment plates for 25 days and generated SSCOs containing both regions of chondrocytes expressing SOX9 and neurons expressing TUJ1 (Fig. 6h,i). Similarly, to explore the default developmental potential of HMMR+ cell-derived SSCOs *in vivo*, we transplanted HMMR+ cell aggregates on day 20 into the kidney capsule of NCG mice, and samples were taken out and dissected 7 weeks after transplantation (Fig. 6j). Interestingly, these transplanted tissues seemed more mature than those generated by NMP-derived SSCOs, and contained osteogenic regions expressing COL-1, chondrogenic regions expressing ACAN, and neuronal regions expressing TUJ1 and MAP2, especially motor neurons expressing ChAT (Fig. 6k,i). These results suggested that HMMR+ cells in SSCOs were a new progenitor cell type with abilities of self-renewal and multipotency but were different from NMPs, which may contribute to body axis elongation *in vivo* and represent a potential cell source for the treatment of spine and spinal cord disorders.

**Fig. 6:**
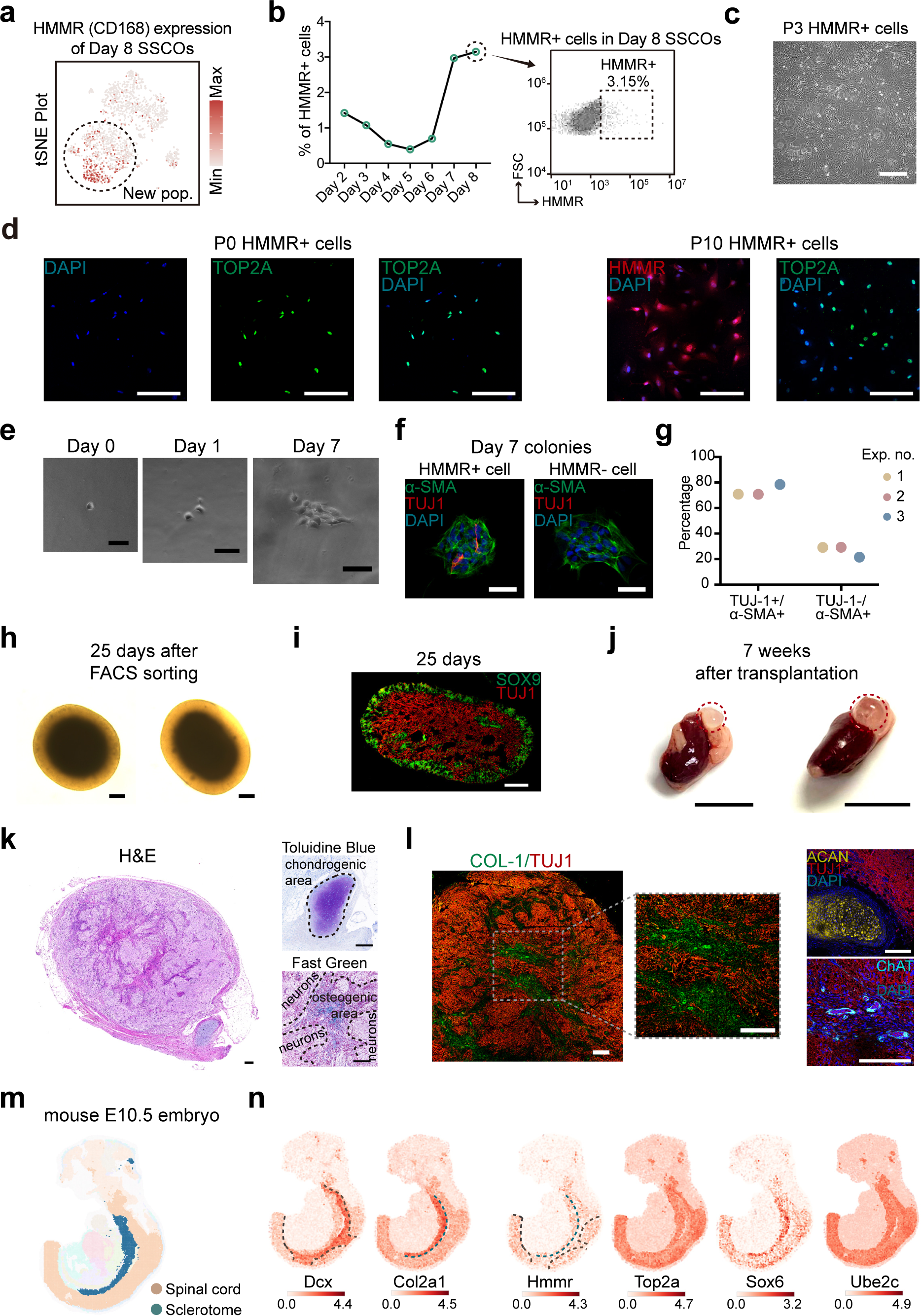
The New HMMR+ Cell Population is bipotent and Can Generate SSCOs. **a,** *t*-SNE plots shows the new population expresses the cell surface marker hyaluronan mediated motility receptor (HMMR) /CD168 significantly. The expression level was scaled from grey (low expression) to red (high expression). **b,** FACS analysis shows the proportion of HMMR+ cells in day8 SSCOs. **c,** Bright-field image of passage 3 HMMR+ cells. Scale bar, 400 μm. **d,** Immunofluorescence analysis shows that both P0 and P10 HMMR+ cells express TOP2A. Scale bar, 200 μm. **e,** Bright-field images of colony formation from individual HMMR+ cells. Scale bar, 75 μm. **f,** Immunofluorescence analysis shows that single HMMR+ cells can generate colonies containing TUJ1+ neurons and α-SMA+ smooth muscle cells, while colonies derived from single HMMR-cells contain only α-SMA+ cells. Scale bar, 50 μm. **g,** Quantitative analysis of the percentage of clones that contain both neural and mesodermal cells (TUJ1+/α-SMA+) or clones that contain only mesodermal cells (TUJ1-/α-SMA+) from three batches. **h,** Bright-field images of organoids derived from HMMR+ cells on day 25. Scale bar, 250 μm. **i,** Immunofluorescence analysis of neural marker (TUJ1) and chondrogenic marker (SOX9) in HMMR+ cell-derived organoids. Scale bar, 200 μm. **j,** Transplants of HMMR+ cell-derived organoids in kidney capsular of NCG mice for 7 weeks. Scale bar, 1 cm. **k,** H&E staining, Toluidine Blue staining and Fast Green staining analysis of sectioned 7-week transplants showing the chondrogenic, osteogenic and neuronal regions. Scale bar, 200 μm. **l,** Immunofluorescence analysis of sectioned 7-week transplants showing the expression of osteogenic marker COL-1, chondrogenic marker ACAN and neuronal markers TUJ1 and ChAT. **m-n,** Mouse organogenesis spatiotemporal transcriptomic atlas (MOSTA) analysis shows the expression and localization of Hmmr+ cells which significantly express *Hmmr, Top2a, Sox6* and *Ube2c* in mouse E10.5 sclerotome and spinal cord.

To verify whether HMMR+ cells exist *in vivo*, we analyzed the human embryo scRNA-seq data from GW2-2.5 whole embryo, GW4-7 spinal cord and GW8-25 spinal cord. The results manifested that HMMR+ cells (HMMR+/TOP2A+/SOX6+) were maintained stably with a ratio about 1-2% in these samples, while NMPs (SOX2+/TBXT+) accounted for 1% of the sequenced cells in GW2-2.5 embryo and almost disappeared at later stages (Extended Data Fig. 6a-c). Similarly, in mouse scRNA-seq data of E6.5-E8.5 embryo (E-MTAB-6967)^51^ and E9.5-E13.5 embryo spinal cord (E-MTAB-7320)^27^, Hmmr+ cells emerged at E7.5 and were still preserved at E13.5. Nonetheless, NMPs that also arose at E7.5 could no longer be detected at E13.5 (Extended Data Fig. 6d-f), which was consistent with the previous report^12, 13^. Next, we checked the distribution and localization of Hmmr+ cells through mouse organogenesis spatiotemporal transcriptomic atlas (MOSTA; https://db.cngb.org/stomics/mosta) ^52^. We identified the Hmmr+ cell zone (Hmmr+/Top2a+/Sox6+/Ube2c+), spinal cord neuron zone (Dcx+) and sclerotome/cartilage zone (Col2a1+). We found that Hmmr+ cells localized at both spinal cord zone and sclerotome zone (Fig. 6m,n and Extended Data Fig. 7a-c). Remarkably, spatial visualization of the Hmmr+ cell zone (Hmmr+) showed distinct but relatively complementary localization pattern especially in E10.5-13.5 spinal cord zone (Dcx+) (Fig. 6m,n and Extended Data Fig. 7a,b). This observation suggests that HMMR+ cells may exist for long term and give rise to sclerotome and spinal cord during body axis elongation as well as later trunk development during pregnancy and postnatal stages after NMP exhausted. In addition, hmmr+ cells were validated through zebrafish embryogenesis spatiotemporal transcriptomic atlas (ZESTA: https://db.cngb.org/stomics/zesta)^53^. Zebrafish hmmr+ cells were discovered in somite and spinal cord region, while NMPs mainly distributed in earlier segmental plate and tail bud region (Extended Data Fig. 7d,e). These results further clarified that this HMMR+ progenitors were conserved between different species and may persist long-term *in vivo* to participate in trunk development after exhaustion of NMPs.

## Discussion

Human organogenesis is a coordinated, orderly, and interdependent process with spatiotemporal tissue patterning under complex but yet undetermined molecular mechanisms. Therefore, *in vitro* recapitulating the morphogenesis of human complex multi-organ structures is necessary and invaluable. The emergence of 3D cultured organoid technology helps to model the development of special organs/tissues. Structurally and functionally integrated hepato-biliary-pancreatic organoids (HBPOs) were developed at the foregut–midgut boundary.^54^ Human neuromuscular organoids (NMOs) from NMPs contained functional neuromuscular junctions which contracted and developed central pattern generator-like neuronal circuits.^55^ A multilineage hiPSC-derived organoid that mimic cardiac and gut development and maturation was also reported.^56^ However, there is still limited research focusing on self-organization of human trunk-like structures and the coordination and interactions of multiple cell lineages during human trunk organogenesis. In the current study, we established the continuous patterning and dynamic morphogenesis of the spine-spinal cord organoids (SSCOs) from hPSC-NMPs through self-organization. SSCOs can further mature and generate complex tissues containing functional motor neurons and chondrocytes/osteoblasts with somitic features *in vitro* and *in vivo*. In addition, the specification of SSCOs could partially imitate the spinal cord development in human GW4-25 embryos according to the scRNA-seq data.

The discovery of NMPs has challenged the traditional paradigm of three germ layers formation during gastrulation and subsequent neural commitment from ectoderm. Indeed, bipotent progenitors with the capacity to differentiate into cells/tissues of two germ layers have been identified, including p75+/HNK1+/SOX10+ neural crest stem cells that can generate peripheral neural lineages and mesenchymal progeny,^57^ and SOX17+/TBXT+/GSC+ mesendodermal progenitors with the capacities to give rise to derivatives of mesoderm and endoderm.^58^ NMPs are a transitional cell population during trunk development *in vivo*^13^ and have been successfully generated from pluripotent stem cells.^59^ Previous reports revealed that PSC-derived NMPs could contribute to posterior neural and mesodermal regions of the embryonic body upon grafting into cultured mouse and chick embryos in xenotransplantation assays.^59, 60^ hPSC-derived NMPs were also used for the generation of human spinal cord organoid.^9^ Nonetheless, whether NMPs could form complex tissues containing both spine and spinal cord *in vitro* has not been addressed, and it still remains unknown about their exact contribution to body axis development and the niche driving the mesodermal and neuroectodermal cell lineage specification. Here, for the first time, we successfully established the spine-spinal cord organoids (SSCOs) from hPSC-NMPs through self-organization. SSCOs recapitulated the morphogenesis of spine/spinal cord and displayed morphological and cellular maturation during the long-term *in vitro* culture, exhibiting an osteogenic trajectory (NMPs-HMMR+ cells-MPs-CPs-CCs-OBs) and a neural trajectory (NMPs-HMMR+ cells-NSCs-NPCs-INs-MNs) just as their *in vivo* developmental dynamics. In addition, the SSCOs showed both rostro-caudal and dorsal-ventral patterning characteristics as described above. Interestingly, our results denoted that HOX gene expression was activated sequentially during the differentiation process and most of HOX paralogs (1-13) were expressed by cells within SSCOs. Therefore, we infer that human NMPs may contribute to hindbrain and all the segments of spinal cord *in vivo*, which was in accordance with the *NKX1-2*-based lineage tracing study.^61, 62^ Moreover, it is reported that NMPs mainly generate ventral tissue in anterior spinal cord and become preponderant in posterior spinal cord by contributing dorsal as well as ventral regions.^10, 61^ Strikingly, Ju-Hyun et al. reported that human NMP-derived spinal cord organoids (hSCOs) did not exhibit obvious dorsoventral patterning in mouse.^9^ Nevertheless, although dorsal-ventral patterning was detected in our SSCO model, dorsal-specific genes were particularly enriched in day 20 SSCOs, while ventral specification was greatly enhanced in day 50 SSCOs. These results somewhat differed from the above literature reports and may be due to the addition of dorsal inducer BMP and ventral inducer SHH agonist in our culture medium, since SHH/BMP pathways are known to antagonize each other during patterning of the dorsal-ventral axis in embryo^31^. The inconsistent findings may reflect the species differences or diverse culture conditions and need further elucidation.

Previous reports showed that NMPs emerged at E7.5 and exhausted at E13.5 when body axis extension ends in mouse embryo, indicating that NMPs did not participate in the trunk development after E13.5.^12^ In our study, NMC-like bipotent HMMR+ cells that derived from NMPs were identified in SSCOs, which did not express *SOX2* or *TBXT* but also possess the ability to generate neural and mesodermal lineages. HMMR usually regulates cell growth and motility and associates cancer progression in multiple tumor types. HMMR is also expressed in the developing nervous system and the proliferative regions of the adult mouse brain, while mutation or knockout of HMMR in mice induces developmental defects of brain.^63^ Here we showed that HMMR+ cells could be propagated *in vitro* and differentiate into motor neurons and somitic mesodermal cells *in vitro* and *in vivo*. Accordingly, we hypothesized that HMMR+ cells represent a potential cell source for the treatment of spine and spinal cord disorders. Furthermore, the *in vivo* scRNA-Seq data from zebrafish, mouse and human embryos demonstrated that this cell population localized at both spinal cord and sclerotome regions, thus NMPs may be truly involved in trunk development in vertebrates. Moreover, HMMR+ cells was first detected at E7.5 and persisted at E15.5 in mouse embryos. These data suggested HMMR+ cell may exist for long term and give rise to sclerotome and spinal cord during body axis elongation as well as later trunk development during pregnancy and postnatal stages. One recent study analyzed the single cell mRNA profiles after spinal cord injury (SCI) and revealed that oligodendrocyte precursor cells (OPCs) proliferated quickly in response to SCI, which play important roles during recovery of SCI.^64^ Intriguingly, we identified a large proportion of Hmmr+ cells concurrently expressing Top2a/Sox6/Ube2c within the OPC cluster (data not shown), suggesting that Hmmr+ cells may also be critically associated with functional recovery of injured spinal cord, which requires to be further clarified.

NMPs may contain heterogeneous cell populations including truly NMPs and lineage-restricted NMCs.^4^ Indeed, although most of the day 2 hPSC-NMPs were positive for SOX2 and TBXT expression, we observed varied staining intensity of these two markers in individual NMPs. It was obvious that some PSC-NMPs with higher expression level of *SOX2* together with lower *TBXT* expression and vice versa (Supplementary Fig. 1c and Supplementary Fig. 3f). NMPs also expressed NKX1-2 and TBX6 (Supplementary Fig. 1h,i). In scRNA-seq data, day 2 NMPs uniformly expressed NMP markers like *CDX1* and *SP5*. Nonetheless, we found that *TBX6* and *NKX1-2* separately labeled different NMP populations (Supplementary Fig. 1h). It has been reported that *Tbx6* plays a key role of presomitic mesoderm (PSM) and somite development, while loss of *TBX6* results in formation of ectopic neural tubes at the expense of PSM.^65, 66^ *NKX1-2* is expressed in the preneural tube and increases rapidly when NMPs are differentiated to preneural cell state.^4^ Based on these data, we hypothesized *TBX6* and *NKX1-2* may play diverse roles during NMP differentiation. Although lineage tracing experiments displayed that NKX1-2+ cells were mainly involved in development of mice neural tube (ectoderm), somites (mesoderm), and other tissues^61^, the exact role of *NKX1-2* have not yet been defined. Thus, we generated *NKX1-2^-/-^*hPSCs by CRISPR/Cas9 technology and induced these mutant cells differentiate to SSCOs. The preliminary data showed nearly complete absence of neural derivatives in organoids, indicating that *NKX1-2* is essential for the posterior spinal cord commitment from NMPs during trunk development (Supplementary Fig. 3). Therefore, the SSCOs model may provide an ideal tool for disease modeling and key gene screening related to congenital spinal diseases. Besides, cell-chat analysis revealed functional cell-cell interaction networks between cell clusters in SSCOs, which may provide exciting opportunities to explore the possible development and maturation mechanisms of different cell lineages during human trunk development in the future.

## Methods

### Mice

All animal experimental procedures were approved by the Animal Ethics Committee of Sun Yat-Sen University, with a project license number of North-F2020-0546QX.

Both male NCG (NOD/ShiLtJGpt-*Prkdc^em26Cd52^Il2rg^em26Cd22^*/Gpt) mice aged 6-8 weeks were purchased from Gempharmatech (Cat# T001475; RRID: IMSR_GPT: T001475) and housed under standard specific-pathogen-free (SPF) conditions with a 12-hour light/dark cycle, a temperature and humidity-controlled facility (20 ± 2 °C; 50 ± 10%) in accordance with the institutional guidelines and ethical regulations, and free access to food and water.

### Human pluripotent stem cell (hPSC) lines and culture conditions

The female human embryonic fibroblast-induced pluripotent stem cell (HEF-iPSC) line^67^ and the male human peripheral blood mononuclear cell-induced pluripotent stem cell (HPBMC-iPSC) line were generated previously. The female human embryonic stem cell line (H9) was obtained from WiCell Research Institute^68^. hPSCs were maintained in feeder-free conditions using mTeSR plus medium (STEMCELL Technologies) on Matrigel (BD Biosciences). Cells were passaged every 3–5 days with ReLeSR (STEMCELL Technologies). All cells used had a normal diploid karyotype.

### Generation and differentiation of hPSC-NMPs

For NMP generation, hPSCs were dissociated into single cells using Accutase (Gibco) at 37°C for 5-8 minutes when hPSCs reached about 70-80% confluency. Cells were counted by Cellometer Auto T4 cell counter (Nexcelom, USA) and replated on Matrigel-(BD Biosciences) coated dishes at a density of 3.5×10^4^ cells/cm^2^ in mTeSR plus medium with 10 μM Y27632 (Sigma-Aldrich) for 24 hours (day 0). Then, cells were cultured in TeSR-E6 medium (STEMCELL Technologies) with 10 μM CHIR99021 (StemRD), 20 ng/ml bFGF (Peprotech), and 5 ng/ml TGFβ1 (Peprotech) for 48 hours (day 0-day 2). The medium was changed every day. Day 2 cells were analyzed for the co-expression of the NMP markers T/BRA and SOX2 by immunofluorescence.

For neural differentiation of hPSC-NMPs, day 2 cells were first cultured in N2B27 medium containing 47.5% DMEM/F12 medium (HyClone), 47.5% Neurobasal medium (Gibco), 1% Penicillin-Streptomycin (HyClone), 1% N2 supplement (Gibco), 2% B27 supplement (Gibco), 1% Glutamax (Gibco), 55 μM 2-mercaptoethanol (Gibco), 1μM RA (Sigma-Aldrich), 10 μM SAG (StemRD), 1 μM SB431542 (StemRD), and 0.5 μM LDN193189 (StemRD) for 4 days to generate NMP-NSCs. Rosette structures were readily observed at day 6. Then day 6 NMP-NSCs were dissociated using Accutase at 37°C for 3-5 minutes, passaged on Matrigel coated dishes in 1:6 ratio and cultured in N2B27 medium supplemented with 1 μM RA, 10 μM SAG, 10 ng/ml GDNF (Peprotech), 10 ng/ml BDNF (Peprotech), 10 ng/ml NT-3 (Peprotech), 0.2 mM AA (Sigma-Aldrich), and 0.5 mM Db-cAMP (Sigma-Aldrich) for 4-6 weeks to generate spinal cord neurons. The medium was changed every 2 days.

For mesenchymal differentiation of hPSC-NMPs, day 2 cells were cultured in NMP induction medium for additional 2 days, and then were dissociated and cultured in MesenCult-ACF Plus Culture medium (STEMCELL Technologies) for 3 weeks to generate NMP-MSCs ^14^. For osteogenic differentiation, NMP-MSCs were cultured in DMEM/Low glucose medium (HyClone) containing 10 mM BGP (Sigma-Aldrich), 0.1 µM DEX (Sigma-Aldrich), 0.2 mM AA, and 10% FBS (HyClone) for 2–3 weeks. For adipogenic differentiation, NMP-MSCs were cultured in DMEM/High glucose medium (HyClone) containing 1 µM DEX, 10 μg/ml Insulin (Sigma-Aldrich), and 0.2 mM IBMX (Sigma-Aldrich), and 10% FBS for 4 weeks. For chondrogenic differentiation, NMP-MSCs were suspended as a sphere in a 15-ml conical tube and cultured in MesenCult-ACF Chondrogenic Differentiation medium (STEMCELL Technologies) for 4 weeks.

For clonal plating experiments, day 2 hPSC-NMPs were dissociated into single cells and replated on Matrigel-coated dishes with a density of 20-50 cells/cm^2^ in N2B27 basal medium with 1 μM RA, 1 μM SB431542, 0.5 μM LDN193189, and 10 μM Y27632 for 8-10 days.

### Generation of spine-spinal cord organoids (SSCOs) in 3D culture

For generation of SSCOs, hPSC-NMPs were dissociated into single cells and cultured in N2B27 medium with 20 ng/ml bFGF, 20 ng/ml EGF (Peprotech), 1μM RA, 10 μM SAG, 1 μM SB431542, and 0.5 μM LDN193189. Cells self-organized into spheroids after 12-24 hours. All spheroids were then transferred into ultra-low 6-well plate (Corning) on the CO2 Incubator Orbital Shaker (C0-06U, SilentShake, China) in 80 rpm and cultured for 5-6 days. For further differentiation, day 8 SSCOs were cultured in N2B27 medium with 1 μM RA, 10 μM SAG, 10 ng/ml GDNF, 10 ng/ml BDNF, 0.2 mM AA, 100 ng/ml TGFβ1, 50 ng/ml BMP4 (Peprotech) for 14-20 days.

To promote the rostrocaudal elongation and maturation of SSCOs, day 20 SSCOs were cultured in 1:1 mixed N2B27 basal medium and MesenCult-ACF Chondrogenic Differentiation medium with 1 μM RA, 10 μM SAG for more than 30 days. Medium was changed every 2 days.

### SSCOs dissociation for scRNA-seq with BD Rhapsody

SSCOs were incubated in Accutase at 37°C for 10-30 minutes (depending on size), and then were mechanically dissociated using a pipette until a single cell suspension was obtained. The single-cell suspension was passed through a 70-μm reversible strainer (STEMCELL Technologies) and pelleted via centrifugation at 500 g for 5 minutes. Cells were resuspended in PBS containing 0.01% BSA (Calbiochem) to give a final concentration of 10^6^ cells/180 μl with adding extra 20 μl tag.

For multiplexing, samples were labeled using the BD Human Single Cell Sample Multiplexing Kit (BD Biosciences, 633781) at room temperature for 20 minutes (or on ice for 30 minutes), pooled and washed with 0.01% BSA-PBS twice for library preparation. Single cells were captured and libraries prepared using the BD Rhapsody system with the BD Rhapsody cDNA Kit (Cat. No. 633773) and Whole Transcriptome Analysis (WTA) Amplification Kit (BD Biosciences, 633801). The WTA and Sample Tag libraries were amplified and purified according to the manufacturer’s protocol (https://www.bdbiosciences.com/content/dam/bdb/marketing-documents/BD_Rhapsody_System_mRNA_Whole_Transcriptome_Analysis_and_Sample_Tag_Library_Preparation_Protocol.pdf). The libraries were pooled and then delivered to the medical inspection laboratories (Annoroad Gene Technology, China) for sequence by Novaseq.

### Analysis of scRNA-seq data

Raw sequencing data were processed with the BD Rhapsody Complete Analysis Pipeline on the Seven Bridges Genomics cloud platform, which resulted in de-multiplexed counts matrices of gene expression in single cells. The R-package Seurat (version 4.1.1) was used for downstream analyses, including quality control, data normalization, data scaling and visualization. Cells that expressed fewer than 200 genes, more than 6,000 unique molecular identifiers, more than 25% of reads assigned to mitochondrial genes or definitive multiplets with two distinct sample tags were filtered out of the analysis. The final dataset contained 11,172 cells. A principal component analysis was used for dimension reduction with a dimension value of 15 determined by the JackStrawPlot function. The top 2,000 variable genes were selected and used together with dimensional information for clustering. Unsupervised clustering was performed, and *t*-SNE for dimension reduction plots were generated. To discover mixed cell clusters, 1893 cells which belongs to day 8 were selected and reclustered, same data processing procedure were performed and a final *t*-SNE for dimension reduction plots were generated with the parameter dims=1: 20.

For cellular cross-talking analysis of clusters in SSCOs, CellChat (version 1.6.0) was performed following the guidelines at: https://github.com/sqjin/CellChat. All the parameter in each function were using the default. In details, the strength of each cell clusters (incoming or outgoing), total numbers of each cell cluster interaction, ligand-receptor pair and outgoing/incoming signaling pattern were checked step by step. Cells which belong to day 20 and day 50 were analyzed independently.

For pseudotime trajectory analysis of SSCOs differentiation, Monocle (version 2.22.0) were used to calculate the cell trajectory, the parameter used were followed at http://cole-trapnell-lab.github.io/monocle-release/docs/ In details, each samples were undergone 6 steps: estimateSizeFactors, estimateDispersions, differentialGeneTest, setOrderingFilter, reduceDimension, orderCells and with no parameter modified.

### Integration of scRNA-seq data

Reciprocal principal component analysis (RPCA) based integration could effectively detect a state-specific cell cluster and runs significantly faster on large datasets. This method was used for our datasets integration. Before the integration, each public datasets were processed follow their description: GW2-2.5 whole embryo (E-MTAB-9388) ^47^, GW4-7 spinal cord (GEO: GSE171892) ^48^, and GW8-25 spinal cord (GEO: GSE136719) ^49^.

Each data was then combined and integrated through Seurat (version 4.0.5) followed the guidelines at: https://satijalab.org/seurat/articles/integration_rpca.html. NMP with different development stages (day 8, day 20 and day 50) were merged with these public scRNA data independently. Differential expression gene analysis on each overlapped cell clusters (e.g., cluster1 in Day 8 merge, cluster 4 and cluster 6 in day 20 merge and cluster 11 in day 50 merge) Function FindAllMarkers embedded in Seurat (version 4.0.5) to find out useful information that mark these overlapping states.

### RNA isolation, reverse transcription, and quantitative PCR analysis

Total RNA of cells or organoids were extracted using TRIzol Reagent (RNAzol) according to the manufacturer’s instructions. RNA yield was determined by using the NanoDrop ND-1000 spectrophotometer (NanoDrop Technologies, USA). Total RNA (1 μg) was converted to cDNA using a NovoScript Plus All-in-one 1st Strand cDNA Synthesis Super Mix kit (Novoprotein). Quantitative real-time PCR (qPCR) analysis was performed using a SYBR Green Mix qPCR kit (Roche) and the LightCycler 480 Detection System (Roche Diagnostics, USA). The expression levels were normalized to those of glyceraldehyde-3-phosphate dehydrogenase (*GAPDH*), changes in gene expression were calculated as fold changes using the delta-delta Ct method, and standard deviations were calculated and plotted using Prism 8 software (GraphPad). Primer details were listed in Supplementary Table 1.

### Immunofluorescence analysis

For immunofluorescence staining, cells were fixed with 4% PFA (PHYGENE) at room temperature for 15 min and washed 3 times with PBS. Then cells were permeabilized with 0.3% Triton X-100 (Sigma-Aldrich) in PBS and incubated overnight at 4 °C with primary antibody or isotype control antibody diluted in PBS containing 10% donkey serum (Jackson) or goat serum (Cell Signaling Technology). Secondary antibodies (1:1000 dilution) were incubated for 1-2 hours at room temperature. Samples were counterstained with 4′,6-diamidino-2-phenylindole (DAPI) (Sigma-Aldrich) and mounted with mounting medium (DAKO).

For immunofluorescence staining of organoids and tissues, samples were first fixed in 4% PFA (PHYGENE) at room temperature for 30 minutes to 4 hours (depending on size), washed with PBS for 3 times, and then left overnight in 30% sucrose solution for dehydration. Organoids and tissues then were embedded with Tissue-Tek O.C.T. Compound (SAKURA) and prepared into frozen sections with cryostat (CM1950, Leica, Germany). Immunofluorescence staining protocol of organoids and tissues was the same as described above.

All confocal images were captured by the confocal laser-scanning microscope (LSM 780, Zeiss, Germany) and the high-speed confocal imaging system (Dragonfly CR-DFLY-202 2540, Andor, UK), and were processed with the corresponding softwares ZEN Blue (Zeiss) and Imaris (Bitplane).

### Histological analysis with whole slide imaging

For histological analysis of SSCOs and tissues, samples were fixed with 4% PFA (PHYGENE) at 4 °C for more than 24 hours. Tissues then were processed for dehydration and wax leaching. The wax-soaked tissue then was embedded and prepared into paraffin sections with pathology slicer (RM2016, Leica, Germany).

Prepared paraffin sections were first dewaxed and used for Hematoxylin and Eosin (H&E) staining (Servicebio), Toluidine Blue staining (Servicebio) and Safranin O-Fast Green staining (Servicebio). All dehydration, dewax, and histological staining were followed the standardized protocol from Servicebio website (https://www.servicebio.cn). All whole slide images of histological staining were captured by the diagnostic scanner (Pannoramic 250 FLASH, 3DHISTECH, Hungary) with the corresponding Pannoramic Scanner software (3DHISTECH) and CaseViewer software (3DHISTECH).

### Calcium imaging of mature neurons in SSCOs

For calcium imaging of mature neurons in SSCOs, SSCOs at day 50 were attached on Matrigel-coated dishes for 2 weeks. Attached cells were washed with Krebs-Henseleit Buffer (adjusting the pH to 7.4 with NaOH) containing 130 mM NaCl, 3 mM KCl, 2.5 mM CaCl, 0.6 mM MgCl2, 10 mM HEPES, 10 mM Glucose, 1.2 mM NaHCO3. Then cells were incubated in Krebs-Henseleit Buffer with 5 μM Fluo-4AM, 2.5 mM Probenecid, and 0.1% Pluronic F-127 (all from Thermo Fisher) at 37 °C for 15 minutes and washed with Krebs-Henseleit Buffer for 3 times. Changes in fluorescence intensity of active cells were recorded using high-throughput live cell imaging system for 10 minutes (BioTek-lionheart FX, BioTek, USA). Fluorescent image stacks were processed by Fiji software ^69^ and plotted using Prism 8 software (GraphPad).

### Electrophysiological recordings of mature neurons in SSCOs

For electrophysiological recordings of mature neurons in SSCOs, artificial cerebrospinal fluid containing 124 mM NaCl, 5 mM KCl, 1.25 mM NaH2PO4, 1 mM MgCl2, 2 mM CaCl2, 26 mM NaHCO3, and 10 mM dextrose gasses with 95% O2/5% CO2 was prepared. Cells in SSCOs were visualized using an upright microscope (Olympus corp, Japan) with infrared differential interference contrast optics and water immersion ×40 objective. Activities from spinal neurons were recorded using MultiClamp700B amplifier and BNC-2090 digitizer (Natinal Inc, USA) in borosilicate glass pipettes with resistances of 3–5 MΩ after filling with intracellular solution containing 10 mM KCl, 10 mM Na-phosphocreatine, 0.2 mM EGTA, 4 mM MgATP, 0.5 mM Na2GTP,10 mM HEPES, 130 mM K-gluconate,10 mM L-glutamic acid, 0.5 mM DNDS, and 0.1-0.2% biocytin, (adjusting the pH to 7.25-7,45 and the osmolarity to 300 mOsm). All the recordings were performed at 35°C. Voltage signals were filtered at 2.8 kHz using a Bessel filter and digitized at 10 kHz. Data were analyzed using the igor software (Wavemetrics inc).

### Subcapsular kidney transplantation of SSCOs in NCG mice

SSCOs from NMPs at day 30 or SSCOs from HMMR+ cells at day 25 were transplanted into the subcapsule of the kidney in immune-deficient NCG mice (age: 6–8 weeks) from Gempharmatech. The incision of kidney subcapsule was plugged using dealmed surgical sponges. Transplanted samples were collected after 1-2 months.

### Fluorescence-activated cell sorting for HMMR+ cells

For Fluorescence-activated cell (FACS) sorting, SSCOs at day 8 were dissociated to single cells by treatment with Accutase at 37°C for 10-15 minutes. After washing with PBS, the single cells were incubated with PE-conjugated HMMR/CD168 antibody (Novus) at room temperature for 15 minutes. The digested cells were filtered through a 70-μm cell strainer. Cell sorting was performed in a high speed cell sorter (MoFlo Astrios EQs, Beckman Coulter, USA) and analyzed with FlowJo software (Flow Jo).

### Maintenance and differentiation of HMMR+ cells

For maintenance of HMMR+ cells after FACS sorting, cells were plated on Matrigel-coated dishes and cultured in N2B27 medium with 20 ng/ml bFGF and 20 ng/ml EGF. Cells were passaged in 1:3 ratio every 3–5 days with Accutase. The method for clonal differentiation of single HMMR+ cells was similar to that of hPSC-NMPs.

For generation of SSCOs, HMMR+ cells (10,000 cells/well) were plated on an round-bottom ultra-low 96-well plate (Corning) in N2B27 medium with 1 μM RA, 10 μM SAG, 10 ng/ml GDNF, 10 ng/ml BDNF, 0.2 mM AA, 100 ng/ml TGFβ1, 50 ng/ml BMP4 for 15 days, then were transferred into ultra-low 6-well plate on the shaker and cultured in 1:1 mixed N2B27 medium and MesenCult-ACF Chondrogenic Differentiation medium with 1 μM RA, 10 μM SAG for more than 10 days.

### Statistical analysis and reproducibility

All experiments shown were biologically replicated at least three times. All results are shown as the mean ± standard deviation (SD) from at least three independent experiments. Unpaired two-tailed Student’s t tests were performed when two groups of samples were compared. All the p values were calculated using GraphPad PRISM 8 with the following significance: n.s. p > 0.05; * p < 0.05; ** p < 0.01; *** p < 0.001; **** p < 0.0001. Statistical details for each experiment can be found in the Fig. and the legends.

### Data availability

The SSCOs single cell data generated this paper were deposited in the NGDC database: https://ngdc.cncb.ac.cn/gsa-human/s/C30vbi4t, with the accession number HRA003724.

## Acknowledgements

This work was supported by the National Key Research and Development Program of China (2022YFA1104100, 2021YFA1100603), the National Natural Science Foundation of China (82270566, 32130046, 81970474), the Key Research and Development Program of Guangdong Province (2019B020236002), the Pioneering Talents Project of Guangzhou Development Zone (2021-L029), and the Key Scientific and Technological Program of Guangzhou City (202206060003).

## Author information

These authors contributed equally: Dairui Li, Yuanchen Ma, Weijun Huang.

## Authors and Affiliations

**Center for Stem Cell Biology and Tissue Engineering, Key Laboratory for Stem Cells and Tissue Engineering, Ministry of Education, National-Local Joint Engineering Research Center for Stem Cells and Regenerative Medicine, Zhongshan School of Medicine, Sun Yat-sen University, Guangzhou, Guangdong, China**

Dairui Li, Yuanchen Ma, Weijun Huang, Chuanfeng Xiong, Qi Zhao, Bin Wang, Shanshan Huang, Yili Wei, Junhua Chen, Xiyu Zhang, Lan Wei, Wenjin Ye, Qiumin Chen, Andy Peng Xiang & Weiqiang Li

**Center of Gastrointestinal Surgery, The First Affiliated Hospital, Sun Yat-sen University, Guangzhou, Guangdong, China**

Yuanchen Ma

**Department of Hepatic Surgery and Liver Transplantation Center of the Third Affiliated Hospital, Organ Transplantation Institute, Sun Yat-sen University, Guangzhou, Guangdong, China**

Xiaoping Li

**Department of Obstetrics and Gynecology, The Third Affiliated Hospital, Guangzhou Medical University, Guangzhou, Guangdong, China**

Huanyao Liu

**Department of Cardiology, The Eighth Affiliated Hospital, Sun Yat-sen University, Shenzhen, Guangdong, China**

Xingqiang Lai

**Department of Spine Surgery, The Third Affiliated Hospital, Sun Yat-sen University, Guangzhou, Guangdong, China**

Limin Rong

## Contributions

Conceptualization, W.L., A.P.X., and L.R.; Methodology, D.L.; Investigation, and Validation, D.L., C.X., Q.Z., X.L., Y.W., J.C., Q.C., X.Z., B.W., and L.W.; Software and Data Curation, Y.M., W.H., X.L., H.L., and W.Y.; Writing – Original Draft, D.L.; Writing – Review & Editing, Y.M., W.H., W.L., and A.P.X.; Funding Acquisition and Supervision, W.L. and A.P.X.

## Corresponding author

Correspondence to Weiqiang Li, Andy Peng Xiang, and Limin Rong.

## Ethics declarations

Competing interests

The authors declare no competing interests.

## Extended Data

**Extended Data Fig. 1.**
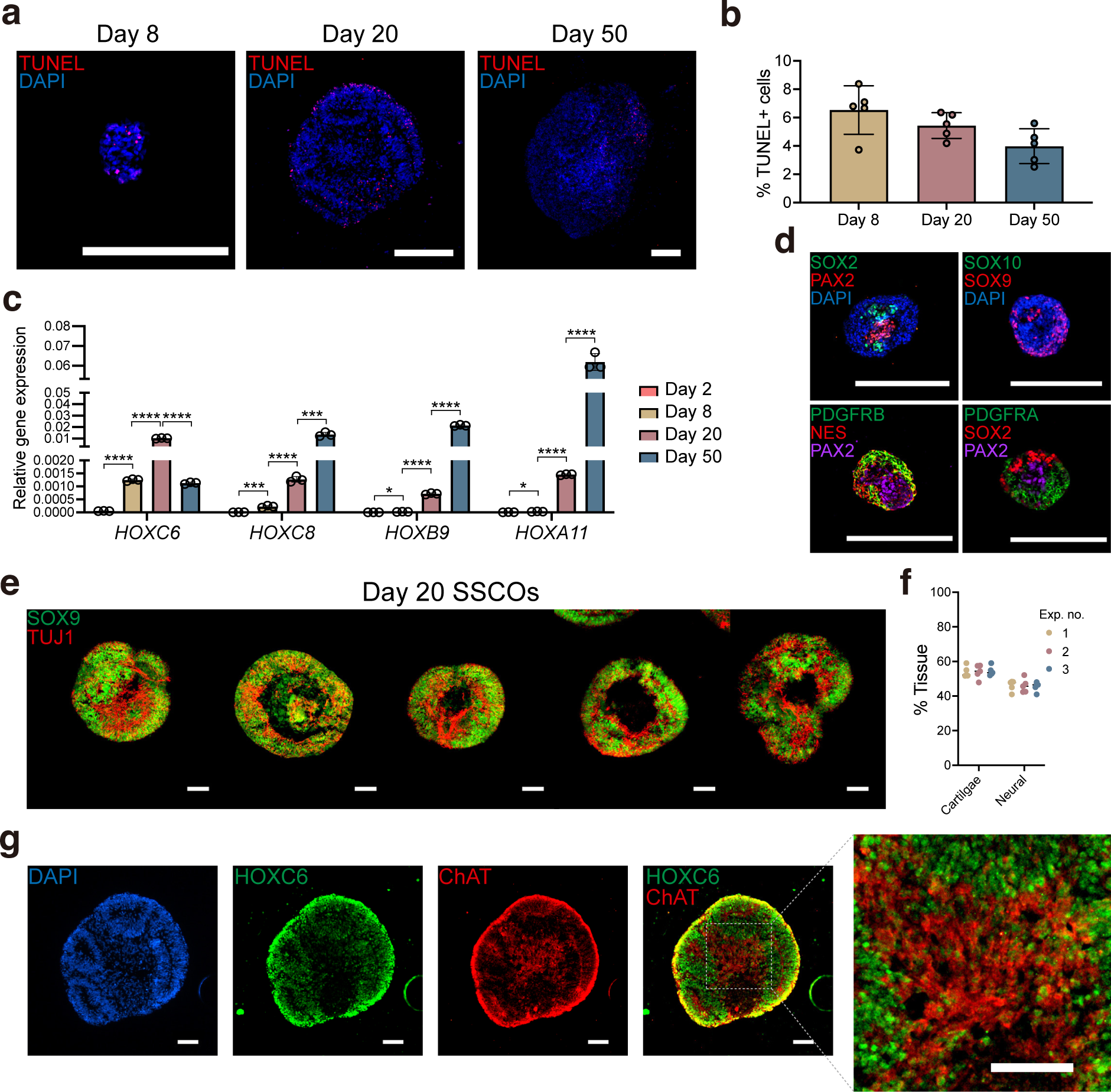
SSCOs were Successfully Generated from Different hPSC Lines. Related to Fig.1, Fig. 2, Fig. 3 and Fig. 4. **a,** Apoptosis cells in SSCOs were detected by TUNEL assay and observed by fluorescence microscopy. Scale bar, 250μm. **b,** Quantitative analysis of the percentage of TUNEL+/DAPI+ cells in SSCOs. **c,** Detection of HOX expression profiles in SSCOs with qRT-PCR. **d,** Immunofluorescence analysis of neural/mesodermal markers in sectioned day 8 SSCOs. Scale bar, 250 μm. **e,** Immunofluorescence staining of day 20 SSCOs from one batch using anti-SOX9 and anti-TUJ1 antibodies. Scale bar: 100μm. **f,** Quantitative analysis of percentage of neural and cartilage tissue area in day 20 SSCOs from 3 batches.

**Extended Data Fig. 2.**
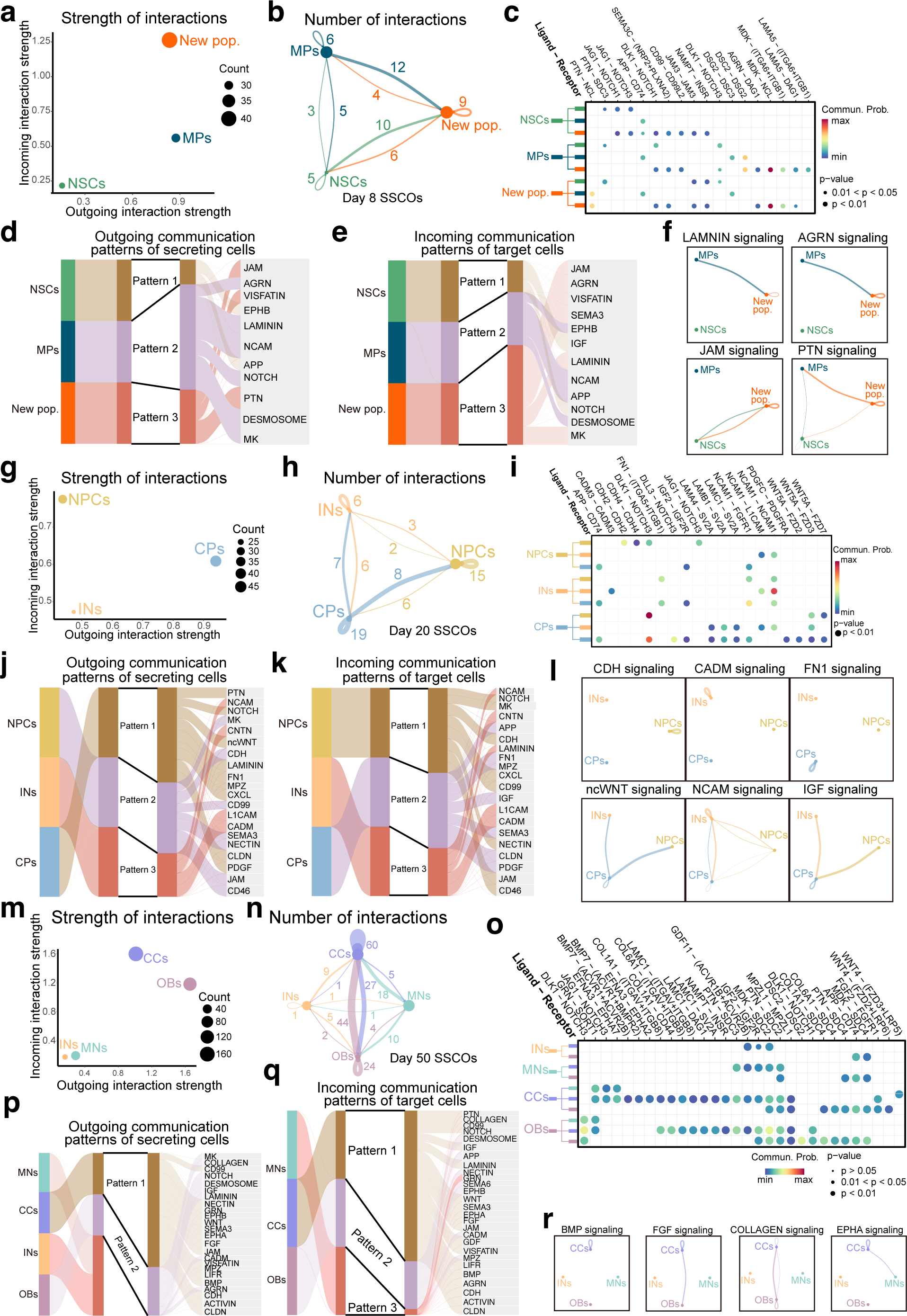
CellChat Analysis of day 8, 20, and 50 SSCOs. Related to Fig. 2, Fig. 3 and Fig. 4. **a,** Total incoming and outgoing interaction weights across three cell populations in day 8 SSCOs. The dot sizes represent the count of ligand-receptor pairs in each cluster. **b,** The number of significant ligand-receptor pairs between any pair of two cell populations. The width of lines is proportional to the indicated number of ligand-receptor pairs. **c,** Selected significant ligand-receptor pairs that contribute to the signaling among new pop., MPs and NSCs. The dot color and size represent the calculated communication probability and p-values, respectively. **d-e,** The outgoing and incoming communication patterns among cells clusters, hierarchical clustering indicated that these cell clusters were separated into three patterns. **f,** Inferred signaling networks between three cell clusters. Circle sizes are proportional to the number of cells in each cell group and line width represents the communication probability. **g,** Total incoming and outgoing interaction weights across three cell populations in day 20 SSCOs. The dot sizes represent the count of ligand-receptor pairs in each cluster. **h,** The number of significant ligand-receptor pairs between any pair of two cell populations. The width of lines is proportional to the indicated number of ligand-receptor pairs. **i,** Selected significant ligand-receptor pairs that contribute to the signaling between mesodermal derivatives represented by CPs and neural derivatives including NSCs and INs. The dot color and size represent the calculated communication probability and p-values, respectively. **j-k,** The outgoing and incoming communication patterns among cells clusters, hierarchical clustering indicated that these cell clusters were separated into three patterns. **l,** Inferred signaling networks between three cell clusters. Circle sizes are proportional to the number of cells in each cell group and line width represents the communication probability. **m,** Total incoming and outgoing interaction weights across four cell populations in day 50 SSCOs. The dot sizes represent the count of ligand-receptor pairs in each cluster. **n,** The number of significant ligand-receptor pairs between any pair of two cell populations. The width of lines is proportional to the indicated number of ligand-receptor pairs. **o,** Selected significant ligand-receptor pairs that contribute to the signaling within mesodermal derivatives including CCs and OBs and neural derivatives including INs and MNs. The dot color and size represent the calculated communication probability and p-values, respectively. **p-q,** The outgoing and incoming communication patterns among cells clusters, hierarchical clustering indicated that these cell clusters were separated into three patterns. **r,** Inferred signaling networks between four cell populations. Circle sizes are proportional to the number of cells in each cell group and line width represents the communication probability.

**Extended Data Fig. 3.**
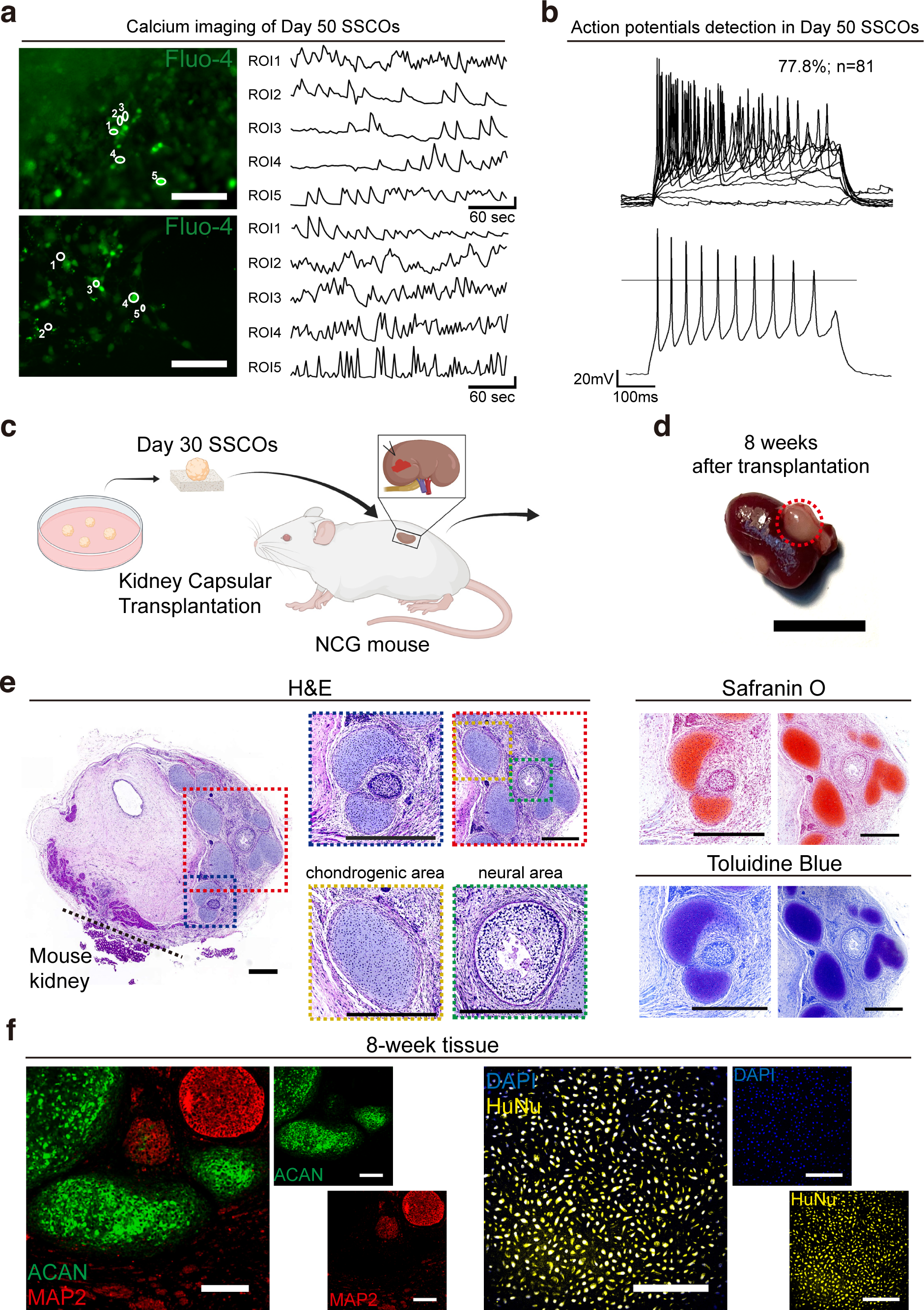
Mature Functional Derivatives Can be Generated in SSCOs *In Vitro* and *In Vivo*. Related to Fig. 3 and Fig. 4. **a,** Calcium imaging of selected neurons in day 50 SSCOs showing continuous flashing green fluorescence by fluo-4 dye (left panel). Scale bar, 100 μm. Line plot showing variation of fluorescence intensities over time for each selected neuron (right panel). **b,** Plot of action potential detected for selected neurons in day 50 SSCOs showing continuously activating action potential (n = 81). **c,** Schematic of kidney capsule transplantation of day 30 SSCOs into NCG mice. **d,** Transplants (circled by red dashed) were detected at the graft site after transplantation for 8 weeks. Scale bar, 1 cm. **e,** H&E staining, safranin O staining and toluidine blue analysis of sectioned 8-week transplants showing the chondrogenic region (yellow dashed frame) and the neural region (green dashed frame). Scale bar, 500 μm. **f,** Immunofluorescence analysis of sectioned 4-week transplants showing the expression of chondrogenic marker ACAN and neuronal marker MAP2 in separate regions and cells in the section are human nuclei (HuNu) positive. Scale bar, 150 μm.

**Extended Data Fig. 4.**
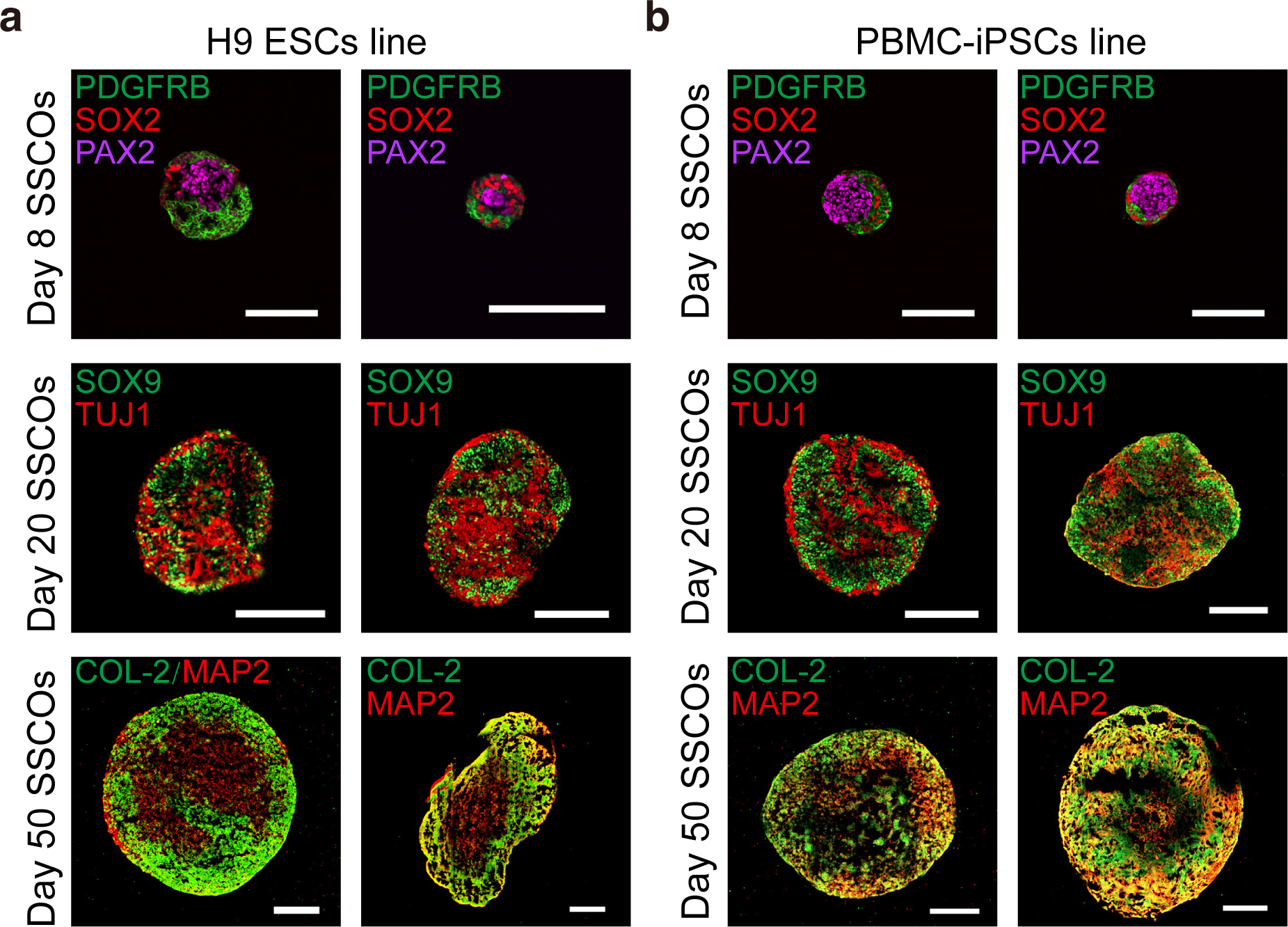
SSCOs were Successfully Generated from Different hPSC Lines. Related to Figure 2, Figure 3 and Figure 4. **a-b,** Immunofluorescence analysis of sectioned day 8, day 20 and day 50 SSCOs showing chondrogenic and neural regions in representative SSCOs at each stage generated from the H9-ESCs line (a), PBMC-iPSCs line (b). Scale bar, 250 μm.

**Extended Data Fig. 5.**
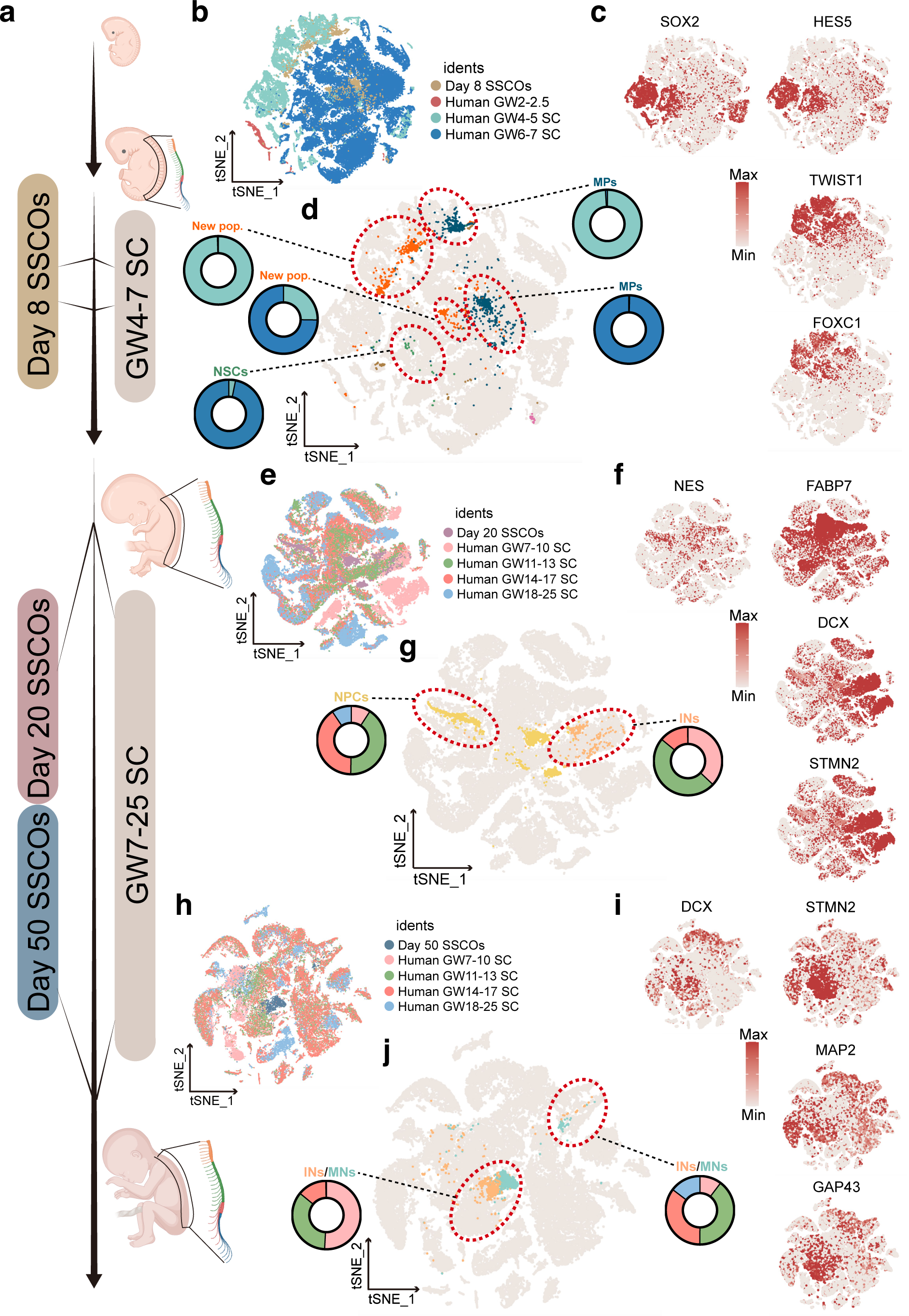
SSCOs Model Human GW4-25 Spinal Cord Development. Related to Fig.2, Fig. 3 and Fig. 4. **a,** Schematic of comparative transcriptome analysis of SSCOs at different stages with whole human embryo (GW2-2.5) and human embryo spinal cord (GW4-7, GW7-25) at the single-cell level. **b,** *t*-SNE plot showing integrative results among day 8 SSCOs, human embryo (GW2-2.5), human embryo spinal cord (GW4-5, GW6-7). Colored by their identity. **c,** *t*-SNE plots showing the individual gene expression levels and distribution of representative marker genes of day 8 SSCOs. SOX2 and HES5 expression indicated the NSC cluster, and TWIST1 and FOXC1 expression indicated the mesoderm cluster. The colors ranging from grey to red indicate low to high relative gene expression levels. **d,** *t*-SNE plot of the identified clusters of day 8 SSCOs corresponding in *in vivo* human data (grey). The pie plot approximate each dotted line was the proportion statistics of each in vivo data, e.g. GW2-2.5 (red), GW4-5 (light blue), GW6-7 (dark blue). **e,** *t*-SNE plot showing integrative results among day 20 SSCOs, human embryo spinal cord (GW7-10, GW11-13, GW14-17, GW18-25). Colored by their identity. **f,** *t*-SNE plots showing the individual gene expression levels and distribution of representative marker genes of day 20 SSCOs. FABP7 expression indicated the NPC cluster, and DCX/STMN2 expression indicated the IN cluster. The colors ranging from grey to red indicate low to high relative gene expression levels. **g,** *t*-SNE plot of the identified clusters of day 20 SSCOs corresponding in *in vivo* human data (grey). The pie plot approximate each dotted line was the proportion statistics of each in vivo data, e.g. GW7-10 (light pink), GW11-GW13 (green), GW14-17 (dark pink) and GW18-25 (blue). h, *t*-SNE plot showing integrative results among day 50 SSCOs, human embryo spinal cord (GW7-10, GW11-13, GW14-17, GW18-25). Colored by their identity. **i,** *t*-SNE plots showing the individual gene expression levels and distribution of representative marker genes of day 50 SSCOs. DCX, STMN2 expression indicated the IN cluster, and MAP2 expression indicated the MN cluster. The colors ranging from grey to red indicate low to high relative gene expression levels. **j,** *t*-SNE plot of the identified clusters of day 50 SSCOs corresponding in *in vivo* human data (grey). The pie plot approximate each dotted line was the proportion statistics of each in vivo data, e.g. GW7-10 (light pink), GW11-GW13 (green), GW14-17 (dark pink) and GW18-25 (blue).

**Extended Data Fig. 6.**
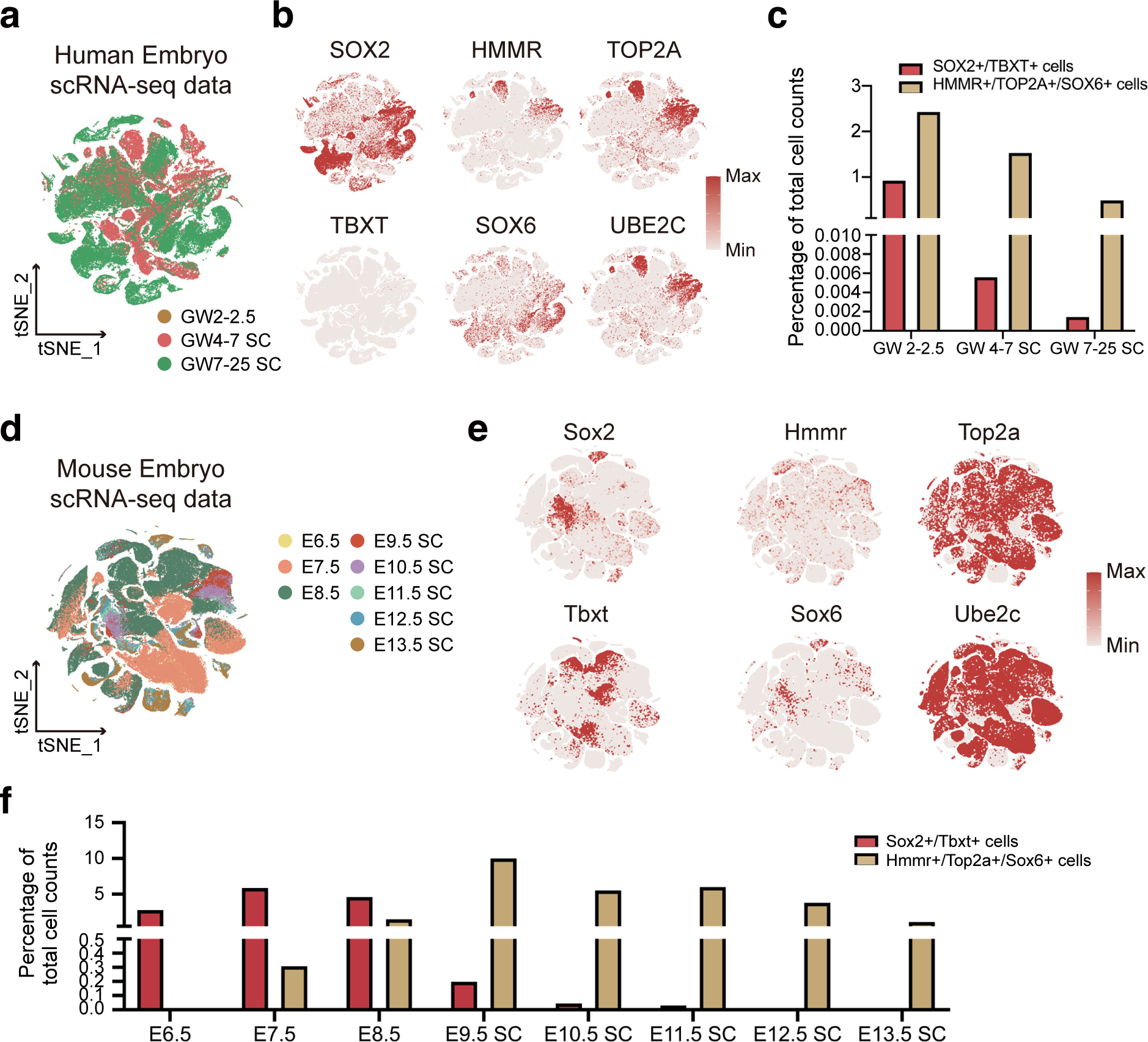
A New HMMR+ Cell Population is Conservative in Human and Mouse embryos. Related to Fig. 5 and Fig. 6. **a,** *t*-SNE plot of scRNA-seq profiles (points) of human embryo (GW2-2.5), human embryo spinal cord (GW4-7) and human embryo spinal cord (GW7-25), colored with three developmental stages. **b,** *t*-SNE plots showing the individual gene expression levels and distribution of NMPs (co-express SOX2 and TBXT) and the HMMR^+^ population (co-express HMMR, TOP2A, SOX6 and UBE2C) in human embryo. The colors ranging from grey to red indicate low to high relative gene expression levels. **c,** Bar chart shows the percentage change of NMPs and HMMR+ cells at different developmental stages of human embryo. **d,** *t*-SNE plot of scRNA-seq profiles (points) of mouse E6.5-8.5 embryo and E9.5-E13.5 embryo spinal cord, colored with their development stage. **e,** *t*-SNE plots showing the individual gene expression levels and distribution of NMPs (co-express Sox2 and Tbxt), and the Hmmr^+^ population (co-express Hmmr, Top2a, Sox6 and Ube2c) in mouse embryo. The colors ranging from grey to red indicate low to high relative gene expression levels. **f,** Bar chart shows the percentage change of NMPs and Hmmr+ cells at different developmental stages of mouse embryo.

**Extended Data Fig. 7.**
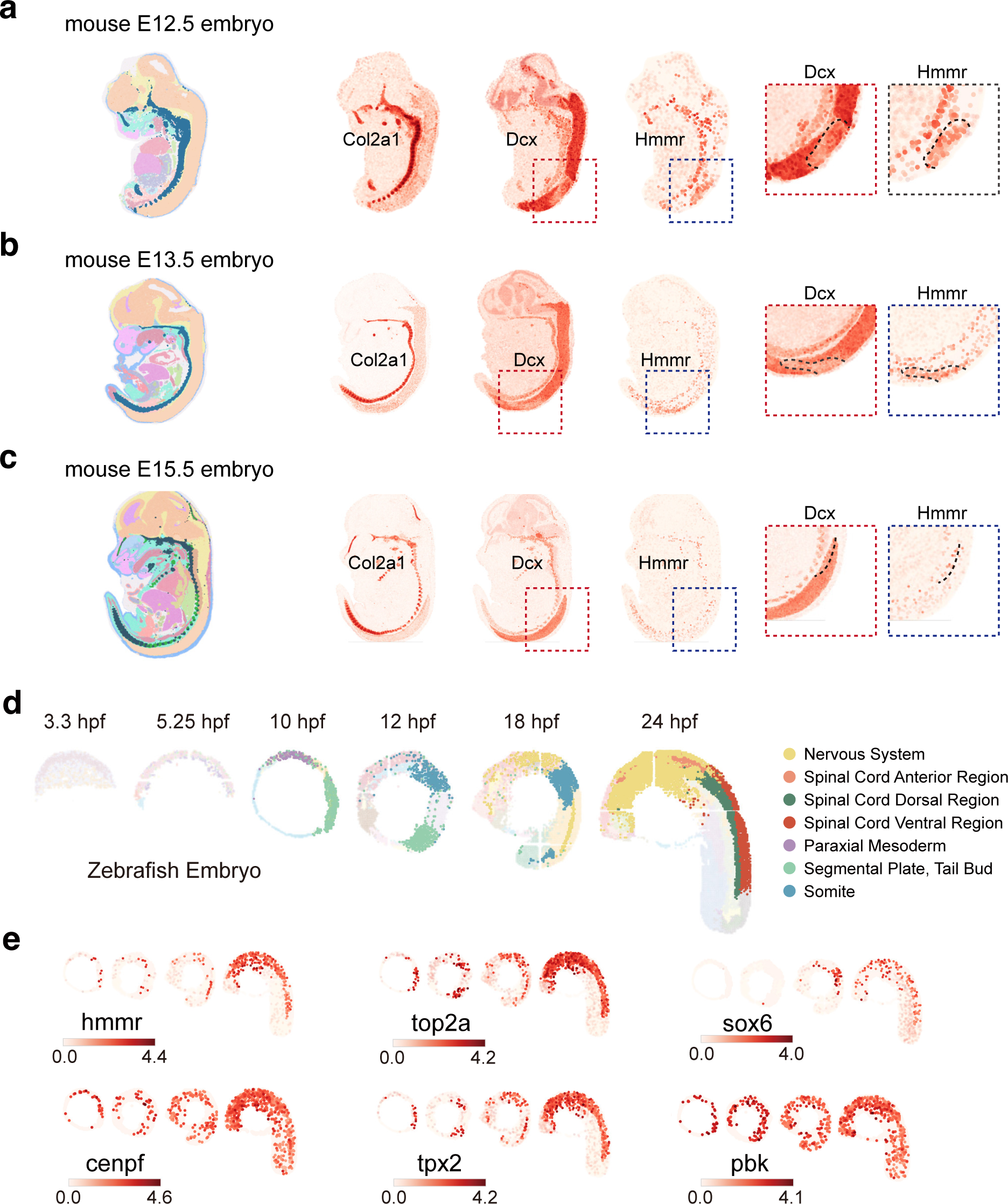
Spatial distribution of Hmmr+ Cells in Mouse Embryo and Zebrafish Embryo. Related to Fig. 5 and Fig. 6. **a-c**, Mouse organogenesis spatiotemporal transcriptomic atlas (MOSTA) analysis showing the location of Hmmr+ cells in spinal cord region (light brown), and sclerotome (dark blue) or cartilage region (light green) were labeled in mouse E11.5-E15.5 embryo (left panel). Markers were used to label spinal cord neurons (Dcx), sclerotome and cartilage (Col2a1), and Hmmr+ cells (Hmmr, Top2a and Sox6) (right panel). **d-e,** Zebrafish embryogenesis spatiotemporal transcriptomic Atlas (ZESTA) analysis showing the location of hmmr+ cells, which significantly express hmmr, top2a, sox6, cenpf, tpx2 and pbk in zebrafish embryonic nervous system, spinal cord (anterior, dorsal and ventral region), paraxial mesoderm, segmental plate and somite. The colors ranging from light pink to red indicate low to high relative gene expression levels.

