## Supplementary figures, tables and videos for "Modeling Human Spine-Spinal Cord Organogenesis by hPSC-Derived Neuromesodermal Progenitors"

**Supplementary Information**

Contents

1. Supplementary Fig. 1 | hPSCs Generate NMPs Efficiently.

2. Supplementary Fig. 2 | hPSC-NMPs Differentiate to Ectodermal and Mesodermal Lineages.

3. Supplementary Fig. 3 | *NKX1-2* is Essential for the Posterior Spinal Cord Commitment from NMPs during Trunk Development.

4. Supplementary Table 1 | Primers Used for qPCR.

5. Supplementary Video 1 | Calcium Imaging of Mature Functional Neurons in SSCOs by Fluo-4 Dye.

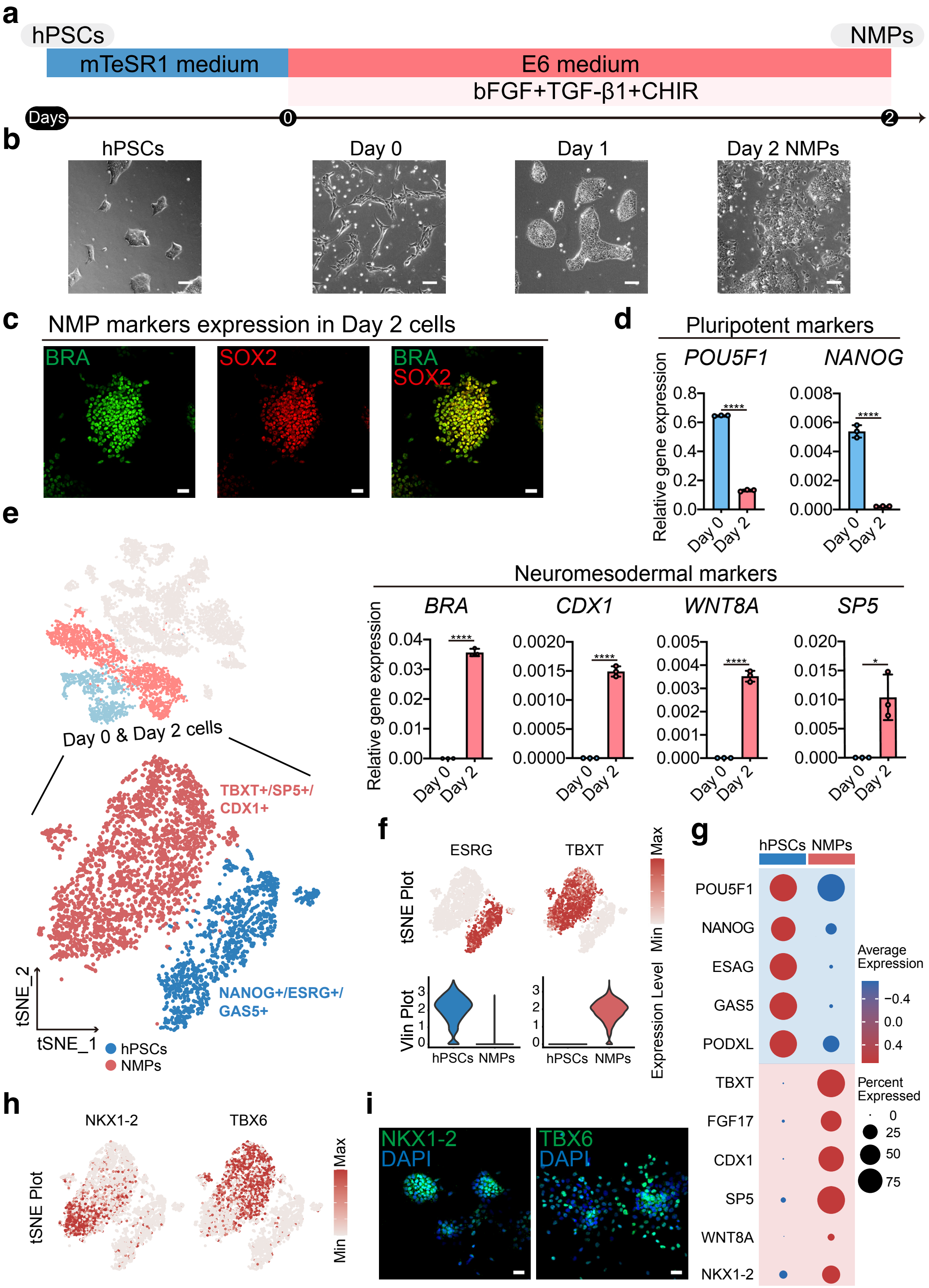


**Supplementary Fig. 1 | hPSCs Generate NMPs Efficiently. Related to Fig. 1**

**a,** Schematic of the strategy used to generate NMPs from hPSCs by activation of FGF, WNT and TGFβ signalings.

**b,** Bright-field images of cell morphology during differentiation process **(a)**. Scale bar, 100 μm.

**c,** Immunofluorescence analysis of day 2 NMPs showing the expression of the NMP markers SOX2, T/BRA, NKX1-2, and TBX6. Scale bar, 40 μm.

**d,** qPCR showing the gene expression of pluripotent markers and NMP markers in day 2 NMPs compared with day 0 hPSCs. Data are represented as mean ± SD. Unpaired two-tailed t test is used for comparison of two groups. *$p < 0.05$, **$p < 0.01$, ***$p < 0.001$, ****$p < 0.0001$.

**e,** *t*-SNE plot of scRNA-seq profiles showing hPSCs colored in blue and NMPs colored in red.

**f,** Gene expression of representative markers among hPSCs and NMPs in *t*-SNE plot (upper panel) and violin plot (lower panel), in *t*-SNE plot each genes distribution and relative expression was scaled from grey (low expression) to red (high expression), and in violin plot the y-axis represents the relative expression level of selected genes.

**g,** Dot plot of cluster-enriched genes of of hPSCs and NMPs. The dot color represents the average expression level of selected genes and dot size represents the percent expressed cells in each cluster.

**h,** Feature plot of selected genes (NKX1-2, TBX6).

**i,** Immunofluorescence analysis of day 2 NMPs showing the expression of NKX1-2, and TBX6. Scale bar, 40 μm.

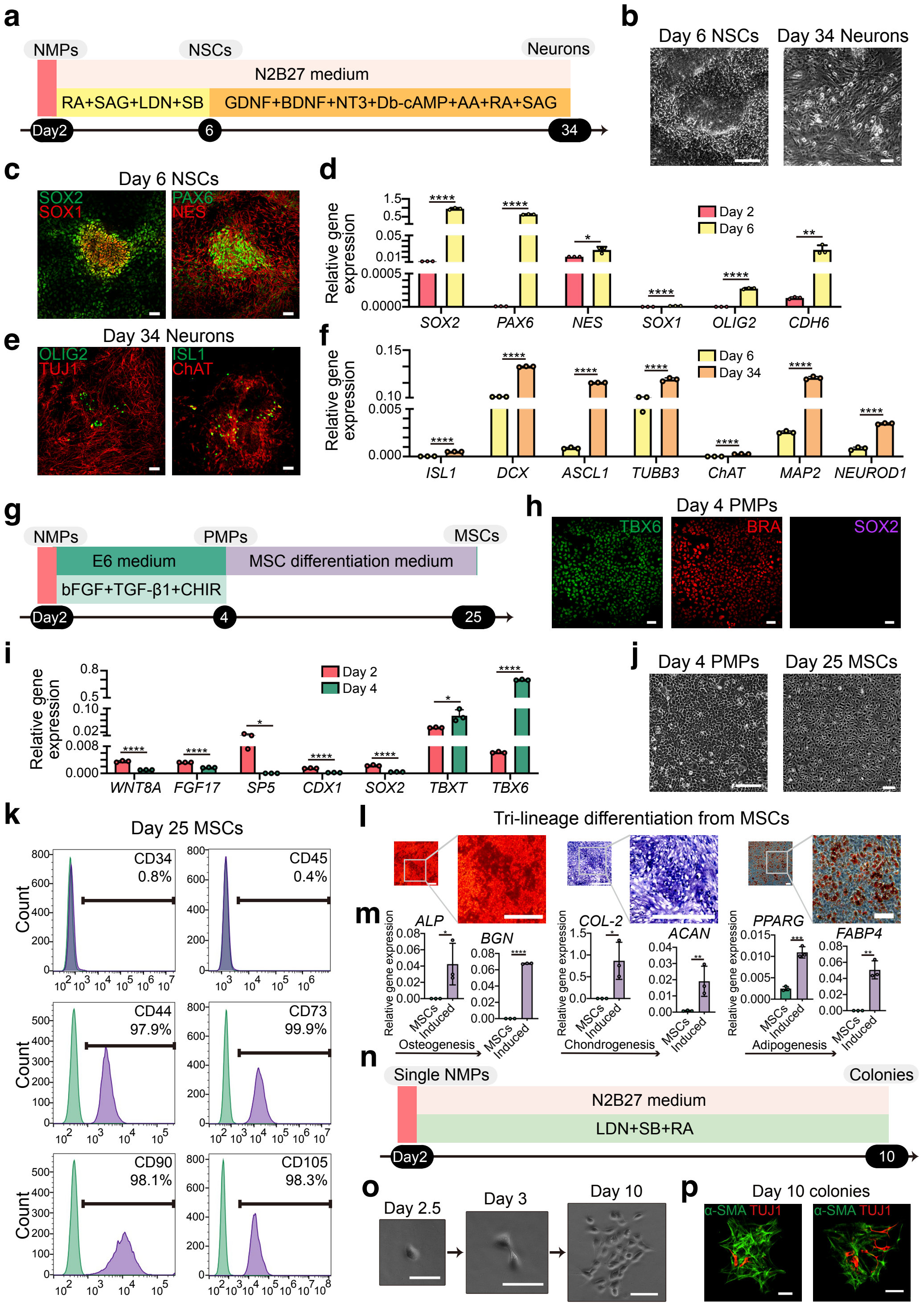


**Supplementary Fig. 2 | hPSC-NMPs Differentiate to Ectodermal and Mesodermal Lineages. Related to Fig. 2, Fig. 3 and Fig. 4**

**a,** Schematic of the strategy used to generate neurons from hPSC-NMPs through NSCs.

**b,** Bright-field images of NSCs showing “neural rosette” structures and neurons during differentiation process **(a)**. Scale bar, 100 μm.

**c,** Immunofluorescence analysis of day 6 cells showing the expression of NSC markers SOX2, SOX1, PAX6 and NES. Scale bar, 50 μm.

**d,** qPCR showing the gene expression of NSC markers in day 6 cells compared with day 2 NMPs. Data are represented as mean ± SD. Unpaired two-tailed t test is used for comparison of two groups. *$p < 0.05$, **$p < 0.01$, ***$p < 0.001$, ****$p < 0.0001$.

**e,** Immunofluorescence analysis of day 34 cells showing the expression of neural markers TUJ1, OLIG2, ISL1 and ChAT. Scale bar, 50 μm.

**f,** qPCR showing the representative gene expression of neural markers in day 34 cells compared with day 6 cells. Data are represented as mean ± SD. Unpaired two-tailed t test is used for comparison of two groups. *$p < 0.05$, **$p < 0.01$, ***$p < 0.001$, ****$p < 0.0001$.

**g,** Schematic of the strategy used to generate mesenchymal stromal cells (MSCs) from hPSC-NMPs through paraxial mesodermal progenitors (PMPs).

**h,** Immunofluorescence analysis of day 4 cells showing the expression of PMP markers T/BRA and TBX6 and the NMP marker SOX2. Scale bar, 50 μm.

**i,** qPCR showing the representative gene expression of NMP markers and PMP markers in day 4 cells compared with day 2 NMPs. Data are represented as mean ± SD. Unpaired two-tailed t test is used for comparison of two groups. *$p < 0.05$, **$p < 0.01$, ***$p < 0.001$, ****$p < 0.0001$.

**j,** Bright-field images of day 4 cells showing the morphology of PMPs and day 25 cells showing the morphology of MSCs. Scale bar, 200 μm.

**k,** FACS analysis showing the expression of cell surface markers in MSCs.

**l,** Bright-field images of Tri-lineage differentiation of NMP-MSCs including osteogenic, chondrogenic, and adipogeneic differentiation on day 25 by Alizarin red S staining (left panel), toluidine blue staining (middle panel), and oil red O staining (right panel) respectively. Scale bar, 200 μm.

**m,** qPCR showing the representative gene expression of osteogenic (*ALP* and *BGN*), chondrogenic (*COL-2* and *ACAN*) and adipogeneic (*PPARG* and *FABP4*) markers in day 25 differentiated cells compared with NMP-MSCs. Data are represented as mean ± SD. Unpaired two-tailed t test is used for comparison of two groups. *$p < 0.05$, **$p < 0.01$, ***$p < 0.001$, ****$p < 0.0001$.

**n,** Schematic of the strategy used to generate colonies from single hPSC-NMPs.

**o,** Bright-field images of colony formation from single hPSC-NMPs. Scale bar, 75 μm.

**p,** Immunofluorescence analysis of colonies **(o)** showing the expression of the neural marker TUJ1 and the mesenchymal marker α-SMA. Scale bar, 75 μm.

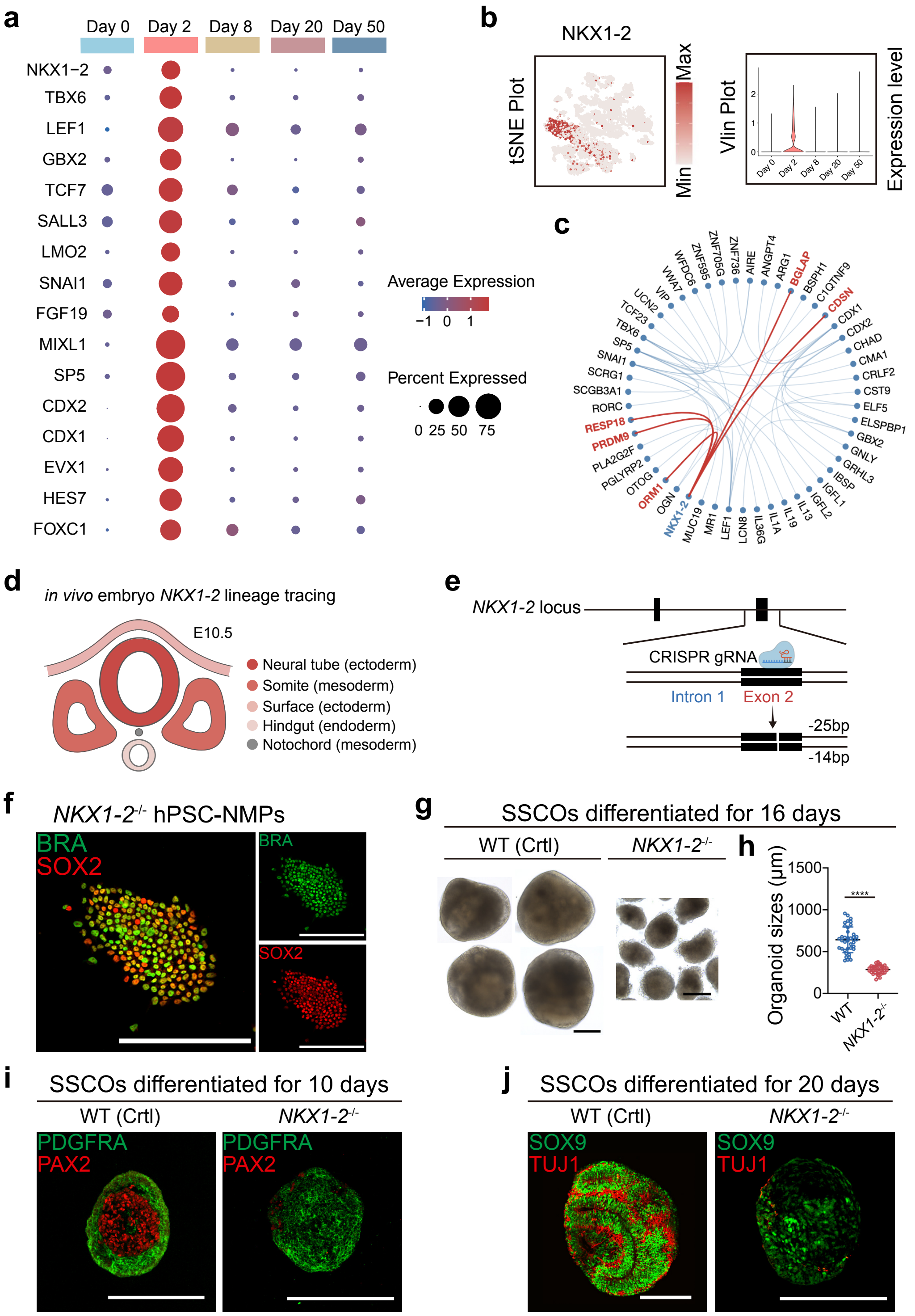


**Supplementary Fig. 3 | *NKX1-2* is Essential for the Posterior Spinal Cord Commitment from NMPs during Trunk Development. Related to Fig. 1, Fig. 2 and Fig. 3**

**a,** Dot plot of day-2-NMP-enriched genes. The colors ranging from blue to red indicate low to high relative gene expression levels.

**b,** *t*-SNE plot and violin plot show that *NKX1-2* was highly enriched in some of hPSC-NMPs.

**c,** Gene interaction analysis shows that *NKX1-2* interacts with both neural genes (*RESP18* and others) and osteogenic genes (*BGLAP* and others). Gene interactions are labeled with red lines.

**d,** Pattern diagram of mouse lineage tracing experiments display that NKX1-2+ cells were mainly involved in development of neural tube (ectoderm), somites (mesoderm), and other tissues (Rodrigo Albors et al., 2018).

**e,** Pattern diagram of construction of *NKX1-2*$^{-/-}$ hPSCs using CRISPR/Cas9 technology.

**f,** Immunofluorescence analysis shows that *NKX1-2*$^{-/-}$ hPSCs can generate NMPs. Scale bar, 250 μm.

**g-h,** Bright-field images of SSCOs from *NKX1-2*$^{-/-}$ hPSC-NMPs and wild-type hPSC-NMPs, and their sizes were calculated and compared. Scale bar, 250 μm. $^{*}p < 0.05$, $^{**}p < 0.01$, $^{***}p < 0.001$, $^{****}p < 0.0001$.

**i-j,** Immunofluorescence analysis of SSCOs shows deficiency in neural commitment of *NKX1-2*$^{-/-}$ hPSC-NMPs compared with wild-type hPSC-NMPs. Scale bar, 250 μm.

**Supplementary Table 1 | Primers Used for qPCR, Related to Fig. 2-4 and Supplementary Fig. 1-2 and Methods**

| Gene | | Primer Sequence (5'-3') |
|---|---|---|
| BRA | forward | TATGAGCCTCGAATCCACATAGT |
| | reverse | CCTCGTTCTGATAAGCAGTCAC |
| TBX6 | forward | AGGCTGTCACGGAGATGAA |
| | reverse | ACAAGTACCAACCCCGCAT |
| SOX2 | forward | GGGAAATGGGAGGGGTGCAAAAGAGG |
| | reverse | TTGCGTGAGTGTGGATGGGATTGGTG |
| PAX6 | forward | TGGGCAGGTATTACGAGACTG |
| | reverse | ACTCCCGCTTATACTGGGCTA |
| MIXL1 | forward | GGCGTCAGAGTGGGAAATCC |
| | reverse | GGCAGGCAGTTCACATCTACC |
| CDH6 | forward | CTGCGACGGATGCAGATGAT |
| | reverse | CCCTGTTTTCTCGATCCATGTTG |
| PAX2 | forward | AGATTCCCAGAGTGGTGTGG |
| | reverse | GGGTATGTCTGTGTGCCTGA |
| FOXC2 | forward | CCTCCTGGTATCTCAACCACA |
| | reverse | GAGGGTCGAGTTCTCAATCCC |
| SOX9 | forward | AGCGAACGCACATCAAGAC |
| | reverse | CTGTAGGCGATCTGTTGGGG |
| NKX3-2 | forward | GATTTCAGGCCTGCTGGGA |
| | reverse | TTTCGCACCCCTTGGTTACA |
| MEOX2 | forward | GGCAAGAGGAAAAGCGACAG |
| | reverse | ATCTCGTATCGCCTCAGTCTG |
| TWIST1 | forward | TTCAAAGAAACAGGGCGTGG |
| | reverse | GCACGACCTCTTGAGAATGC |
| DCX | forward | TCCCGGATGAATGGGTTGC |
| | reverse | GCGTACACAATCCCCTTGAAGTA |
| ISL1 | forward | GCGGAGTGTAATCAGTATTTGGA |
| | reverse | GCATTTGATCCCGTACAACCT |
| NEFL | forward | ATGAGTTCCTTCAGCTACGAGC |
| | reverse | CTGGGCATCAACGATCCAGA |
| IBSP | forward | CACTGGAGCCAATGCAGAAGA |
| | reverse | TGGTGGGGTTGTAGGTTCAAA |
| ACAN | forward | GTGCCTATCAGGACAAGGTCT |
| | reverse | GATGCCTTTCACCACGACTTC |
| MSX1 | forward | AAAGTGGCTGGAAGAGTCCC |
| | reverse | GACACCGATTTCTCTGCGCT |
| TUBB3 | forward | TTCATCTTTGGTCAGAGT |
| | reverse | GCAGGCAGTCGCAGTTTT |
| MAP2 | forward | CGAAGCGCCAATGGATTCC |
| | reverse | TGAACTATCCTTGCAGACACCT |
| LBX1 | forward | GCGGCAGACCCCTAAGAAG |
| | reverse | GGTGATGACTTGCGCGTTG |

| *Continued* | | |
|---|---|---|
| **Gene** | | **Primer Sequence (5'-3')** |
| HB9 | forward | CTCCTACTCGTACCCGCAG |
| | reverse | TTGAAGTCGGGCATCTTAGGC |
| CTSB | forward | AGAGTTATGTTTACCGAGGACCT |
| | reverse | GATGCAGATCCGGTCAGAGA |
| RUNX2 | forward | AGAAGGCACAGACAGAAGCTTGA |
| | reverse | AGGAATGCGCCCTAAATCACT |
| COL-1 | forward | CAGCCGCTTCACCTACAGC |
| | reverse | TTTTGTATTCAATCACTGTCTTGCC |
| ALP | forward | TGAGGGTGTGGCTTACCAG |
| | reverse | GATGGACGTGTAGGCTTTGCT |
| OPN | forward | AGATGGGTCAGGGTTTAGCC |
| | reverse | CATCACCTGTGCCATACCAG |
| POU5F1 | forward | GACAGGGGGAGGGGAGGAGCTAGG |
| | reverse | CTTCCCTCCAACCAGTTGCCCCAAAC |
| NANOG | forward | CAGCCCAGATTCTTCCACCAGTCCC |
| | reverse | CGGAAGGTTCCCAGTCGGGTTCACC |
| CDX1 | forward | GGTGGCAGCGGTAAGACTC |
| | reverse | TGTAACGGCTGTAATGAAACTCC |
| WNT8A | forward | GAACTGCCCTGAAAATGCTCT |
| | reverse | TCGAAGTCACCCATGCTACAG |
| SP5 | forward | GAAACAGTGCTCGGGTTTTC |
| | reverse | TCAACGGGGTCTTCCTGTAG |
| NES | forward | AACAGCGACGGAGGTCTCTA |
| | reverse | TTCTCTTGTCCCGCAGACTT |
| SOX1 | forward | CAGTACAGCCCCATCTCCAAC |
| | reverse | GCGGGCAAGTACATGCTGA |
| OLIG2 | forward | CCAGAGCCCGATGACCTTTTT |
| | reverse | CACTGCCTCCTAGCTTGTCC |
| ASCL1 | forward | CGCGGCCAACAAGAAGATG |
| | reverse | CGACGAGTAGGATGAGACCG |
| NEUROD1 | forward | ATGACCAAATCGTACAGCGAG |
| | reverse | GTTCATGGCTTCGAGGTCGT |
| ChAT | forward | CTCTGGTGTACTCAGCTACAAGG |
| | reverse | CTCAGGCTCCGGCATGATG |
| BGN | forward | GAGACCCTGAATGAACTCCACC |
| | reverse | CTCCCGTTCTCGATCATCCTG |
| PPARG | forward | CGAGAAGGAGAAGCTGT TGG |
| | reverse | TCAGCGGGAAGGACTTTATGTATG |
| FABP4 | forward | ACTGGGCCAGGAATTTGACG |
| | reverse | CTCGTGGAAGTGACGCCTT |

**Supplementary Video 1 | Calcium Imaging of Mature Functional Neurons in SSCOs by Fluo-4 Dye, Related to Extended Data Fig. 3**

Calcium imaging of selected neurons in day 50 SSCOs showing continuous flashing green fluorescence by fluo-4 dye (left panel). Scale bar, 100 μm.
